# Inferring fine-scale rates of mutation and recombination in the coppery titi monkey (*Plecturocebus cupreus*)

**DOI:** 10.64898/2026.01.13.699361

**Authors:** Vivak Soni, Cyril J. Versoza, John W. Terbot, Gabriella J. Spatola, Karen L. Bales, Jeffrey D. Jensen, Susanne P. Pfeifer

**Affiliations:** Center for Evolution and Medicine, School of Life Sciences, Arizona State University, Tempe, AZ, USA; Department of Psychology, University of California, Davis, CA, USA; California National Primate Research Center, Neuroscience and Behavior Division, Davis, CA, USA; Department of Neurobiology, Physiology, and Behavior, University of California, Davis, CA, USA

**Keywords:** primate, Pitheciidae, fine-scale mapping, divergence, mutation, recombination

## Abstract

Despite being a primate of considerable biomedical interest, particularly as a model for social behavior and neurobiology, the evolutionary processes shaping genetic variation in the coppery titi monkey (*Plecturocebus cupreus*) remain largely uncharacterized. Utilizing divergence and polymorphism data together with a recently published high-quality, annotated genome, we here infer the first fine-scale maps of mutation and recombination rates in this platyrrhine. We find a mean genome-wide mutation rate of between 0.93 × 10^-8^ and 1.61 × 10^-8^ per site per generation and a mean genome-wide recombination rate of 0.975 cM/Mb, in line with fine-scale rates estimated in other primates. In addition to providing novel biological insights into the mutation and recombination rates in this emerging model species for behavioral research, these fine-scale maps also improve our understanding of how the processes of mutation and recombination shape genetic variation in the coppery titi monkey genome, and their incorporation into evolutionary models will be a necessary aspect of future downstream inference of other evolutionary processes required to elucidate the genetic factors underlying the phenotypic traits studied in this species.

## INTRODUCTION

Mutation and recombination are important evolutionary processes that shape levels and patterns of genetic diversity in populations. Germline mutations are the ultimate source of novel genetic variation, whilst recombination shuffles this variation into potentially novel haplotypes via crossover and non-crossover events. The rate of input of new mutations, as well as the rate of recombination events, have been shown to vary at every level of measurement: across the Tree of Life, between and within species, and across the genome (for mutation rate variation, see the reviews of Baer et al. 2007; Lynch 2010; Hodgkinson and Eyre-Walker 2011; Pfeifer 2020a; for recombination rate variation, see the reviews of Ritz et al. 2017; Stapley et al. 2017; Johnston 2024).

Both mutation and recombination rate estimation can be performed either via direct observation from pedigrees, or indirectly from sequenced population samples, (though classical disease-incidence approaches have also historically been utilized for mutation rate estimation in humans; Haldane 1932, 1935). The direct estimation of both processes relies on high-throughput genome sequencing of parent-offspring trios or multi-generation pedigrees, counting the number of *de novo* mutations as well as crossover and non-crossover events that have occurred from one generation to the next (see the review of Pfeifer 2020a for an overview, and Pfeifer 2021; Bergeron et al. 2022 for a discussion of the challenges in direct rate estimations). Due to the rarity of both spontaneous mutations and meiotic exchange events in vertebrates, resolution with such direct estimation approaches is relatively coarse, given the small number of generations generally considered (see the review of Clark et al. 2010). Consequently, they provide a genome-wide rate estimate of mutation and recombination, as opposed to a fine-scale map of rate heterogeneities across the genome that is necessary for a variety of applications, including genome-wide association studies and selection scans.

By contrast, indirect mutation and recombination rate estimation are performed on species-level divergence data and population-level polymorphism data, respectively. Central to such indirect mutation rate estimation approaches is the observation that the neutral mutation rate is equal to the neutral divergence rate (Kimura 1968, 1983), with the number of substitutions that accumulate in a lineage being proportional to the per-generation mutation rate. Thus, historically-averaged mutation rates across the divergence time between the target species and an outgroup species can be inferred from phylogenetic sequence data in neutral genomic windows, thereby generating a fine-scale genomic map of mutation rate heterogeneity. However, there is often great uncertainty in both the generation time of a species, and the divergence times between the species under investigation. Estimated mutation rates are therefore given across a range of likely generation and divergence times in order to span this uncertainty. Instead of divergence data, indirect recombination rate estimation approaches rely on population-level data of unrelated individuals for the inference of historical recombination rates from observed patterns of linkage disequilibrium (LD; see the reviews of Stumpf and McVean 2003; Peñalba and Wolf 2020), again utilizing neutral genomic windows to generate fine-scale maps across the genome. These inferred rates are necessarily sex-averaged, and one must account for other population genetic processes that can alter LD (e.g., selection and population history; Dapper and Payseur 2018; Samuk and Noor 2022) and thus potentially confound recombination rate inference. To limit the impact of such confounding factors on the indirect inference of both mutation and recombination rates, high-quality genome annotations are necessary to identify regions of the genome that are evolving neutrally; additionally, a well-fitting demographic model is necessary in the case of recombination rate inference (Johri et al. 2020, 2022).

Although it is common practice to model mutation and recombination as a single mean genome-wide rate, accounting for rate heterogeneity across the genome is critical when performing downstream inference of other population genetic processes. For example, inference of population history, the distribution of fitness effects, and of recent positive and balancing selection can all be confounded when heterogeneity in mutation and recombination rates are unaccounted for (Soni et al. 2023, 2024; Soni and Jensen 2024; and see Dapper and Payseur 2018; Samek and Noor 2022; Ghafoor et al. 2023) due to the interactions between evolutionary processes. For example, Hill-Robertson effects (Hill and Robertson 1966; Felsenstein 1974) are expected to be modulated by the locus-specific recombination environment (Maynard Smith and Haigh 1974; Begun and Aquadro 1992; Charlesworth et al. 1993; and see Charlesworth and Jensen 2021, 2022).

Initial estimates of mutation and recombination rates in primates were largely focused upon humans and other great apes (e.g., Kong et al. 2002; Auton et al. 2012; Stevison et al. 2016). More recent studies have performed inference of these processes in a number of other catarrhines, as well as in species of biomedical importance and extinction risk (e.g., Pfeifer 2020b; Xue et al. 2020; Wall et al. 2022; Versoza, Weiss et al. 2024; Soni, Versoza et al. 2025a, 2025b; Versoza et al. 2025; Versoza, Lloret-Villas et al. 2025; Terbot et al. 2025; Versoza et al. 2026a, 2026b; and see the reviews of Tran and Pfeifer 2018; Soni et al. 2025). The recent publication of a chromosome-level genome assembly that includes protein-coding gene annotations for the coppery titi monkey, *Plecturocebus cupreus* (Pfeifer et al. 2024), provides opportunities to explore mutation and recombination landscapes in a, as of yet, genomically under-researched platyrrhine despite the considerable biomedical interest in the species (e.g., Bales et al. 2007; Lau et al. 2024; and see Bales et al. 2021). In this study, we utilize a combination of patterns of variation within and divergence between coppery titi monkeys and humans, as well as gene-level annotations to mask directly selected genomic regions, in order to indirectly infer fine-scale mutation and recombination rate maps across the coppery titi monkey genome. In addition to providing novel biological insights into mutation and recombination rates in this emerging model species for behavioral and neurobiological research, these fine-scale maps of observed rate heterogeneity will also prove important for the future downstream inference of other evolutionary processes, necessary to elucidate the genetic factors underlying the phenotypic traits studied in this species.

## MATERIALS AND METHODS

### Animal subjects

This study was performed in compliance with all regulations regarding the care and use of captive primates, including the NIH Guidelines for the Care and Use of Animals and the American Society of Primatologists’ Guidelines for the Ethical Treatment of Nonhuman Primates. Procedures were approved by the UC-Davis Institutional Animal Care and Use Committee (protocol 22523).

### Species-level divergence data

To obtain species-level divergence, we needed to identify neutral substitutions between the genomes of the coppery titi monkey and humans. To do so, we first replaced the outdated, scaffold-level *P. cupreus* genome assembly in the 447-way multiple species alignment (available from: https://cglgenomics.ucsc.edu/november-2023-nature-zoonomia-with-expanded-primates-alignment/; Zoonomia Consortium 2020; Kuderna et al. 2024) with the chromosome-level NCBI reference genome for the species (GenBank assembly: GCA_040437455.1; Pfeifer et al. 2024). To this end, we performed the following steps:

1. We removed the scaffold-level *P. cupreus* genome assembly from the 447-way multiple species alignment using the *halRemoveGenome* function implemented in HAL v.2.2 (Hickey et al. 2013).
2. We extracted the neighboring reconstructed ancestral genomes (i.e., PrimatesAnc157 and PrimatesAnc189) from the 447-way multiple species alignment using HAL’s *hal2fasta* function.
3. We aligned the extracted reconstructed ancestral genomes (PrimatesAnc157 and PrimatesAnc189) to the chromosome-level *P. cupreus* genome (PleCup_hybrid) using Cactus v.2.9.2 (Armstrong et al. 2020), maintaining the branch lengths that had been inferred in the 447-way multiple species alignment.
4. We attached this newly generated sub-alignment back into the 447-way multiple species alignment using HAL’s *halReplaceGenome* function.

With the fully annotated *P. cupreus* genome assembly integrated into the 447-way multiple species alignment, we next extracted the sub-alignment containing the genomes for coppery titi monkeys, humans and their reconstructed ancestor (PrimatesAnc003) using Cactus’ *cactus-hal2maf* function. We converted the extracted sub-alignment back to .hal format using HAL’s *maf2hal* function and identified fixed differences between the coppery titi monkey and human genomes using HAL’s *halSnps* function. We focused on point mutations that were on the coppery titi monkey branch by selecting sites where humans and the reconstructed ancestor (PrimatesAnc003) shared the same allele while coppery titi monkeys exhibited a different allele. Given the relatively long divergence time between coppery titi monkeys and humans, we limited our analyses to high-confidence regions in the sub-alignment. Specifically, we excluded sites in regions containing gaps and missing nucleotides (denoted by a “–“ and “N” in the sub-alignments, respectively); additionally, we limited our analyses to regions to which we could confidently map our short-read data by applying a mappability mask to the *P. cupreus* genome assembly, generated using the SNPable pipeline with a read length of 150 bp and a stringency parameter of 1 (https://lh3lh3.users.sourceforge.net/snpable.shtml). Finally, in order to obtain neutral substitutions, we removed sites that are polymorphic in either coppery titi monkeys (see “Population-level polymorphism data”) or humans (using the population-level data of the Yoruban population included in the 1000 Genomes Project; 1000 Genomes Project Consortium 2015) as well as those located within 10 kb of functional regions (based on the protein-coding gene information available for the *P. cupreus* genome assembly [Pfeifer et al. 2024] and the catalogue of regulatory elements constraint across primates [Kuderna et al. 2024]).

### Inferring fine-scale neutral divergence between *P. cupreus* and *H. sapiens*

Based on the species-level divergence data, we inferred fine-scale neutral divergence between *P. cupreus* and *H. sapiens* across genomic windows (with window sizes of 1 kb, 10 kb, 100 kb and 1 Mb, and step sizes of half of the respective window sizes) by dividing the number of neutral substitutions by the number of accessible sites in each genomic window (thereby requiring that ≥ 10% of a window is accessible). Assuming divergence times of 32, 33 and 36 million years between *P. cupreus* and *H. sapiens* (Glazko and Nei 2003), and generation times of 6 and 9 years (Pacifici et al. 2013; Perez et al. 2013), we then calculated the fine-scale neutral divergence rate by dividing by the divergence time in generations.

### Population-level polymorphism data

We obtained population-level polymorphism data from six unrelated coppery titi monkeys paired-end sequenced on an Illumina NovaSeq 6000 to high-coverage. In brief, we removed adapter sequences and trimmed both low-quality and polyG tails using fastp v.0.24.0 (Chen et al. 2018) before aligning the reads to the chromosome-level *P. cupreus* genome (Pfeifer et al. 2024) using the Burrows–Wheeler Aligner v.0.7.15 (Li 2013), with shorter split alignments flagged as secondary using the *-M* option. Prior to variant discovery, we marked duplicated reads using the Genome Analysis Toolkit (GATK) v.4.4.0 *MarkDuplicates* function (van der Auwera and O’Connor 2020) to reduce support from redundant coverage (Pfeifer 2017), and recalibrated the base quality scores of the reads using GATK’s *BaseRecalibrator* and *ApplyBQSR* functions together with a set of high-confidence variants previously obtained in pedigreed individuals (Versoza et al. 2026a). Using these high-quality recalibrated reads (*--minimum-mapping-quality* 40), we called variant and invariant sites (*-ERC* BP_RESOLUTION) separately for each sample using the GATK *HaplotypeCaller*, disabling PCR indel modeling (*-pcr-indel-model* NONE) in accordance with developer guidance for PCR-free library design. We subsequently combined individual gVCFs (*CombineGVCFs*) and jointly genotyped across samples (*GenotypeGVCFs*, with the *-all-sites* flag enabled). In order to obtain high-quality variants, we limited the call set to regions in which all samples exhibited at least half, but no more than double, the genome-wide average coverage, located farther than 5 bp away from the nearest insertion/deletion, and re-genotyped biallelic single nucleotide polymorphisms (SNPs) with complete genotype information across all samples using the graph-based genotyper Graphtyper v.2.7.2 (Eggertsson et al. 2017) to improve genotyping accuracy. Owing to reduced sequencing depth on the X and Y chromosomes, we restricted this re-genotyped dataset to autosomal variants that passed all built-in sample-and site-level quality filters and that were located in mappable regions of the genome (as determined by the SNPable mappability mask; see “Species-level divergence data”). Finally, we phased the resulting dataset using WhatsHap v.2.8 (Martin et al. 2023) and limited recombination rate inference to fully phased SNPs found within alignments ≥ 10 kb in length.

### Inferring fine-scale recombination rates in *P. cupreus*

Based on the phased population-level polymorphism data, we inferred fine-scale recombination rates in *P. cupreus* using two widely applied LD-based approaches: LDhat v.2.2 (McVean et al. 2002, 2004; Auton and McVean 2007) and LDhelmet v.1.10 (Chan et al. 2012). To this end, we performed the following steps:

*LDhat*:

1. Using LDhat’s *complete* function, we generated a lookup table for every two-locus haplotype configuration in our sample of six diploids (via the argument *-n* 12), based on a maximum population-scaled recombination rate ρ of 100 (*-rhomax* 100), a grid size of 201 (*-n_pts* 201), and the empirically estimated 𝜃 of 0.0043/site (*-theta* 0.0043).
2. Based on this lookup table, we generated region-based estimates of ρ (in window sizes of 4,000 SNPs with a 200 SNP overlap) using LDhat’s *interval* function, with a block penalty of 5 (*-bpen* 5), 60 million iterations (*-its* 60000000), and sampling every 40,000 iterations (*-samp* 40000).
3. To ensure convergence, we discarded the MCMC burn-in, using the argument *-burn* 500 implemented in LDhat’s *stat* function.
4. To obtain chromosome-scale estimates of ρ, we combined the region-based estimates of ρ at the midpoint of the overlapping windows, whilst masking localized peaks with ρ > 100 between adjacent SNPs together with their neighboring 50 SNPs (masking a total of 6,327 SNPs across 120 regions) in order to minimize the impact of artificial LD breaks generated by genome assembly errors (see Auton et al. 2012; Pfeifer 2020b).
5. Finally, we calculated the per-generation recombination rate, *r*. To do so, we calculated the effective population size (*N_e_*) based on the mean empirical value of 𝜃 of 0.0043/site and a mutation rate of 1.07 × 10^-8^/site/generation, and used the resulting value of *N_e_* to calculate *r* from ρ.

### LDhelmet

1. As LDhelmet requires sequence information in .fasta format as input, we converted the population-level polymorphism data using the *consensus* function implemented in BCFtools v.1.14 (Danecek et al. 2021), with the *-s* argument enabled to allow for a multi-sample input and the *-H* 1 and *-H* 2 arguments enabled to obtain the first and second haplotypes, respectively, and subsequently concatenated the resulting files per chromosome.
2. We generated a configuration file using LDhelmet’s *find_confs* function, utilizing a window size of 50 SNPs (*-w* 50).
3. We generated a likelihood lookup table using LDhelmet’s *table_gen* function, based on the empirically estimated 𝜃 of 0.0043/site (*-t* 0.0043) and the default grid values of ρ (*-r* 0.0 0.1 10.0 1.0 100.0).
4. We generated padé coefficients for the sampling step using LDhelmet’s *pade* function, based on the mean empirical value of 𝜃 of 0.0043/site (*-t* 0.0043) and the default number of padé coefficients (*-x* 11).
5. We generated a mutation matrix from our empirical data, following the approach outlined in Chan et al. (2012). In brief, we polarized the coppery titi monkey SNPs by identifying the corresponding positions in the ancestral PrimatesAnc157 genome from the 447-way multiple-species alignment, and counted the number of each mutational type in alignments ≥ 10 kb using BEDTools *nuc* v.2.3.0 (Quinlan and Hall 2010).
6. After this data pre-processing, we inferred recombination rates along each chromosome using LDhelmet’s *rjmcmc* function, based on a window size of 50 SNPs (*-w* 50), block penalties (*-b*) of 5, 10, 20, and 50, and our mutation matrix, with a burn in of 100,000 iterations (*--burn_in* 100000) and 1 million total iterations (*-n* 1000000).
7. Finally, we post-processed the binary data output from the previous step using LDhelmet’s *post_to_text* function, obtaining the mean (*-m*) value of ρ for each window, which was converted into *r* via a calculation of *N_e_* using the empirical 𝜃 of 0.0043/site (as in the LDhat step 5 description).

### Assessing the performance of recombination rate estimators under the inferred populations history of *P. cupreus*

To assess the performance of the two recombination rate estimators utilized, we simulated 10 replicates of a 1 Mb region in msprime v.1.3.2 (Baumdicker et al. 2022) with a constant per-site recombination rate of 10^-8^, under the demographic model of *P. cupreus* inferred by Terbot et al. (2026), sampling six individuals to match our empirical data. For each replicate, we generated a random set of nucleotides to create a reference sequence and then drew from the empirical mutational matrix to assign polymorphisms at positions determined by the simulation. Recombination rates were inferred on each simulated dataset with both LDhat and LDhelmet to assess their performance under the species-specific population history.

To account for differences in performance as well as uncertainties in *N_e_*, we re-scaled the recombination rate estimates obtained with LDhat and LDhelmet to the autosomal genetic map length inferred from pedigreed individuals (2,450 cM; Versoza et al. 2026b) using scaling factors of 1.21 and 0.168, respectively.

### Inferring recombination hotspots

In order to infer recombination hotspots, we ran LDhot v.0.4 (Auton et al. 2014) on the landscape of recombination inferred by LDhat via the following steps:

1. We used LDhot’s *ldhot* function to perform 1,000 simulations with a 1.5 kb window size, a 1 kb step size, and a 50 kb background window centered on the hotspot.
2. We used LDhot’s *ldhot_summary* function to combine significant windows, merging adjacent candidates. For calling hotspots, we used a significance threshold of 0.001, whilst a threshold of 0.01 was used for merging hotspots.
3. Finally, we filtered out spurious hotspots based on the recommendations from both the Great Ape Recombination Project (Stevison et al. 2016) and Brazier and Glémin (2024). Briefly, hotspot candidates with a width larger than 10 kb were removed, as well as those with an intensity below 4, or above 200.

Afterwards, we used FIMO v.5.5.7 (Grant et al. 2011) to check how many of the final hotspots contained the putative PRDM9 binding sequence (CCTGCCTCAGCCTCC) recently identified through computational analyses (Versoza et al. 2026b). To assess statistical significance, we used BEDTools *random* v.2.3.0 (Quinlan and Hall 2010) to randomly draw the same number of coldspot regions from the genomic background.

### Assessing correlations between genomic features

We calculated summary statistics for a variety of genomic features — including nucleotide diversity (based on our population-level polymorphism data), divergence (based on our species-level divergence data), recombination rate (based on our estimates obtained with LDhat), as well as CpG-content, GC-content, gene-content, and repeat-content (based on the species’ genome annotations; Pfeifer et al. 2024) — across the 22 autosomes of the coppery titi monkey genome and calculated partial Kendall’s rank correlations across 1 kb, 10 kb, 100 kb, and 1 Mb windows (requiring a minimum accessibility of 50% in both the population genomic data as well as the 447-way multi-species alignment) using the *kendalltau* package implemented in SciPy v.1.16.3 (Virtanen et al. 2020).

## RESULTS AND DISCUSSION

### Population-level polymorphism and species-level divergence data

To estimate neutral divergence as well as fine-scale rates and patterns of mutation and recombination, we obtained population-level polymorphism data from six captive coppery titi monkeys (three males and three females), sequenced at a depth of 38–57× per individual. Alignment-based variant discovery across the autosomes yielded 6.9 million phased SNPs, with an observed transition-to-transversion ratio of 2.6 (Supplementary Table S1; and see “Materials and Methods” for details). With this population genomic data on hand, we next replaced the outdated, scaffold-level *P. cupreus* genome included in the 447-way multiple species alignment (Zoonomia Consortium 2020; Kuderna et al. 2024) with the fully annotated, chromosome-level genome of Pfeifer et al. (2024). Using this updated multiple sequence alignment, we counted the fixed differences between *P. cupreus* and the *P. cupreus–H. sapiens* reconstructed ancestor, PrimatesAnc003, in high-confidence regions (excluding any gaps and applying a mappability mask to the alignment; see “Materials and Methods” for details). In order to examine neutral genomic data, we masked functional regions as well as 10 kb flanking regions. To obtain substitutions, we removed sites that were observed to be polymorphic in either *P. cupreus* or *H. sapiens*. Neutral divergence was subsequently calculated by counting neutral substitutions across genomic windows of various sizes (1 kb, 10 kb, 100 kb, 1 Mb). Supplementary Figure S1 provides the distribution of neutral divergence for each window size.

### The landscape of mutation in the coppery titi monkey

Based on the mean genome-wide fine-scale divergence estimate of 0.056 across 100 kb windows, we calculated the mutation rate across a range of divergence times between coppery titi monkeys and humans (32, 33, and 36 million years ago [mya]; Glazko and Nei 2003), and coppery titi monkey generation times (6 and 9 years; Pacifici et al. 2013; Perez et al. 2013), given that there is uncertainty in both parameters. Across this range of divergence and generation times, as well as across our window sizes, the mean mutation rate ranged between 0.93 × 10^-8^ and 1.61 × 10^-8^ /site/generation. Supplementary Table S2 summarizes the range of mean mutation rates for different divergence times, generation times, window sizes, and accessibility length thresholds (i.e., the minimum number of sites that must be accessible for a window to be considered when calculating mutation rates), whilst Figure 1a provides density plots of neutral mutation rate estimates and Figure 1b portrays the genome-wide, per-site, per-generation rates across the autosomal coppery titi monkey genome (and see Supplementary Figure S2 for estimates from individual autosomes). These mutation rate estimates are notably higher than those inferred from divergence data in the common marmoset (*Callithrix jacchus*), another platyrrhine of biomedical interest (ranging between 0.25 × 10^-8^ and 0.37 × 10^-8^ /site/generation; Soni, Versoza et al. 2025b); however, they are consistent with those inferred in the great apes (e.g., Venn et al. 2014; Jónsson et al. 2017; Tatsumoto et al. 2017; Besenbacher et al. 2019; and see the reviews of Tran and Pfeifer 2018; Chintalapati and Moorjani 2020) and gray mouse lemurs (1.52 × 10^-8^ /site/generation with a 95% CI of 1.28 × 10^-8^ – 1.78 × 10^-8^ /site/generation; Campbell et al. 2021).

**Figure 1.**
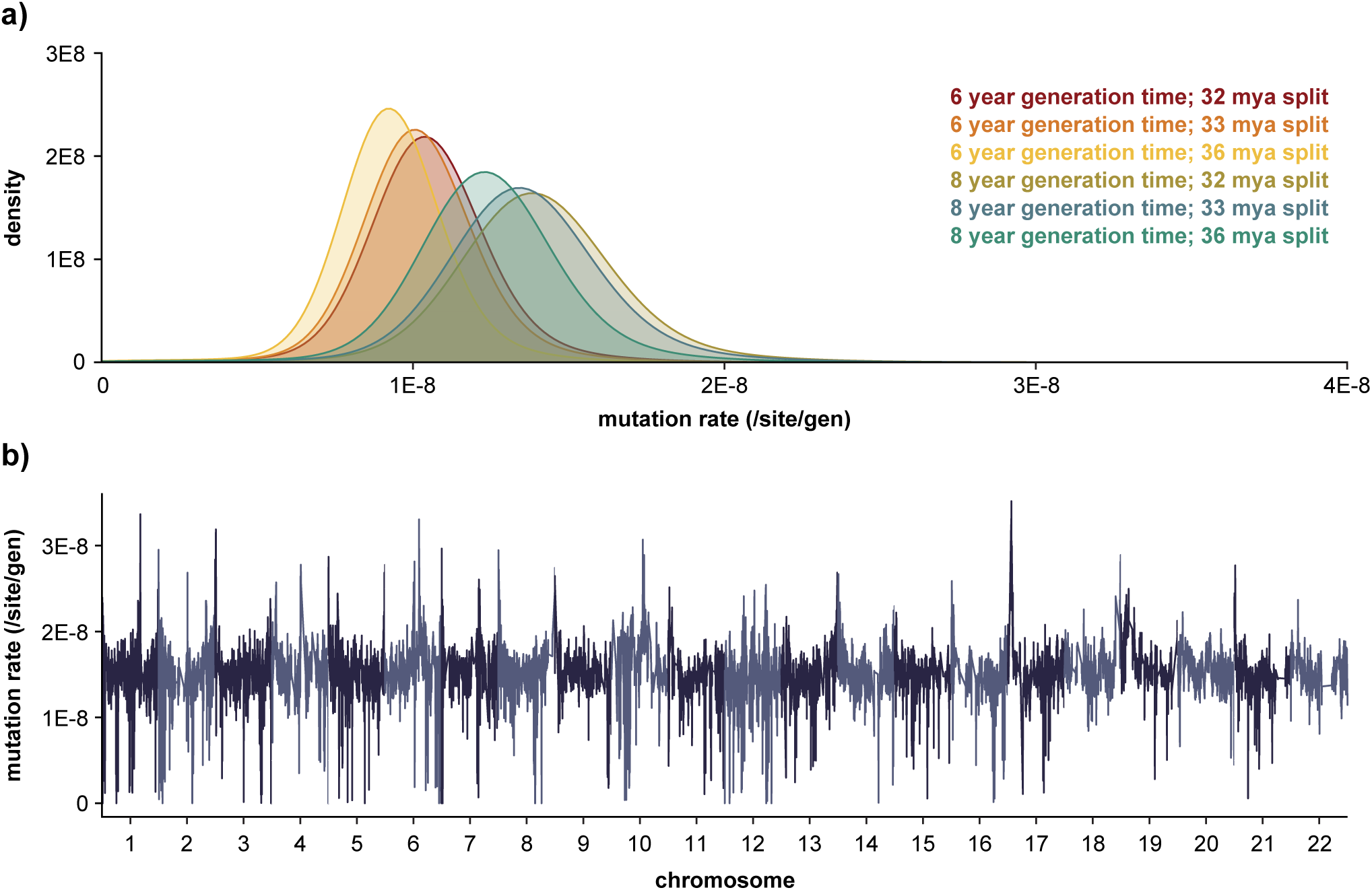
Fine-scale rates of neutral mutation. (a) Density plots of the per-site per-generation mutation rate implied by the neutral divergence for two possible generation times (6 years and 9 years; Pacifici et al. 2013; Perez et al. 2013) and three possible divergence times between the coppery titi monkey (*P. cupreus*) and humans (*H. sapiens*) (32 million years ago [mya], 33 mya, and 36 mya; Glazko and Nei 2003). (b) Genome-wide per-site per-generation neutral mutation rates for genomic windows of size 100 kb, with a 50 kb step size, assuming a *P. cupreus–H. sapiens* divergence time of 33 mya and a generation time of 9 years (and see Supplementary Figure S2 for the heterogeneity in neutral mutation rates across all autosomes). Neutral mutation rates were estimated from the rates of neutral divergence observed between *P. cupreus* and *H. sapiens*.

Helpfully, a recent study utilized pedigree data to provide a genome-wide direct estimation of point mutation rates in *P. cupreus* (Versoza et al. 2026a), allowing us to compare direct and indirect estimates. The authors inferred rates of 0.5 × 10^-8^ /site/generation in individuals born to younger parents and 1.1 × 10^-8^ /site/generation in individuals born to older parents, with an average rate of 0.6 × 10^-8^ /site/generation across the pedigreed individuals. Based on these mutation rate estimates, we calculated divergence times based on our divergence estimate of 0.056 and generation times of 6 and 9 years. Divergence times between coppery titi monkeys and humans ranged from 32 mya (for mutation rates of 1.1 × 10^-8^ /site/generation and generation times of 6 years) to an infeasible 96 mya (for mutation rates of 0.5 × 10^-8^ /site/generation and generation times of 9 years) (Table 1). Although generation times in wild individuals remain elusive, previous studies suggest that titi monkeys reach sexual maturity between 15 months (males) and 32 months (females; Conley et al. 2022), juveniles leave their family group around the age of 2 to 3 years, and adults exhibit life spans of around 20 years in the wild (Zablocki-Thomas et al. 2023) and around 25 years in captivity (de Magalhães and Costa 2009). In captivity, females tend to give birth to their first offspring around the age of 3.7 years (with a range between 2.0 and 6.9 years), and interbirth intervals tend to be around 1.0 to 1.5 years on average (Valeggia et al. 1999; Van Belle et al. 2016). Given previous support for a split time between 32 and 36 mya (Glatzo and Nei 2003), these results thus support a younger generation time together with mutation rate estimates of around (or slightly less than) 1.1 × 10^-8^ /site/generation in wild titi monkey populations. Alternatively, mutation rates higher than 1.1 × 10^-8^ /site/generation would be needed to reconcile an older generation time with our current understanding of primate split times. Taken together, the pedigree-based inference from older parents are highly consistent with our indirect inference provided here, and both are consistent with previous estimates of this split time.

**Table 1.** Inferred *P. cupreus–H. sapiens* divergence times based on the observed mean neutral divergence rate of 0.056 for two different possible generation times (6 years and 9 years; Pacifici et al. 2013; Perez et al. 2013) and three different pedigree-based mutation rate estimates (0.5 × 10^-8^, 0.6 × 10^-8^, and 1.1 × 10^-8^ /site/generation) obtained from parents of differing ages by Versoza et al. (2026a) (shown in orange). Relatedly, the resulting divergence-based mutation rate estimates based on three possible divergence times between the coppery titi monkey and humans (32 million years ago [mya], 33 mya, and 36 mya; Glazko and Nei 2003), and the two possible generation times (shown in blue).

|  |  | pedigree-based mutation rate |  |  | divergence time |  |  |
| --- | --- | --- | --- | --- | --- | --- | --- |
|  |  | 0.5E-08 | 0.6E-08 | 1.1E-08 | 32 mya | 33 mya | 36 mya |
| generation time<br>(years) | 6 | 64 mya | 54 mya | 32 mya | 1.05E-08 | 1.02E-08 | 0.93E-08 |
|  | 9 | 96 mya | 80 mya | 48 mya | 1.58E-08 | 1.53E-08 | 1.40E-08 |

### The landscape of recombination in the coppery titi monkey

Fine-scale estimators of recombination rates are based upon patterns of LD in population-level genomic data, and we inferred the landscape of recombination across the coppery titi monkey genome using two widely applied approaches: LDhat (McVean et al. 2002, 2004; Auton and McVean 2007) and LDhelmet (Chan et al. 2012).

Firstly, to assess the performance of these recombination rate estimators within the context of the specific population history of this species, we simulated a 1 Mb region under the *P. cupreus* demography recently inferred by Terbot et al. (2026). Briefly, the coppery titi monkey population was inferred to have experienced three historical population size changes, including a population expansion ∼131,000 generations ago, with the population increasing from ∼45,000 individuals to almost 2 million, before undergoing a more recent, severe collapse in population size to ∼12,300 individuals occurring 3,160 generations ago. We simulated 10 replicates of this history with a constant recombination rate of 10^-8^ /site/generation. Our simulations demonstrate that both LDhat and LDhelmet underestimate the recombination rate under the *P. cupreus* demographic model (Figure 2). In support of this observation, Dutheil (2024) found via simulation that LDhat underestimates recombination rates in populations that have undergone a recent reduction in population size, as is the case here (note that this study did not investigate the performance of LDhelmet). These results thus again highlight the importance of evaluating the performance of recombination rate estimators within the context of the specific demographic history of the population in question (see the discussion in Johri et al. 2022).

**Figure 2.**
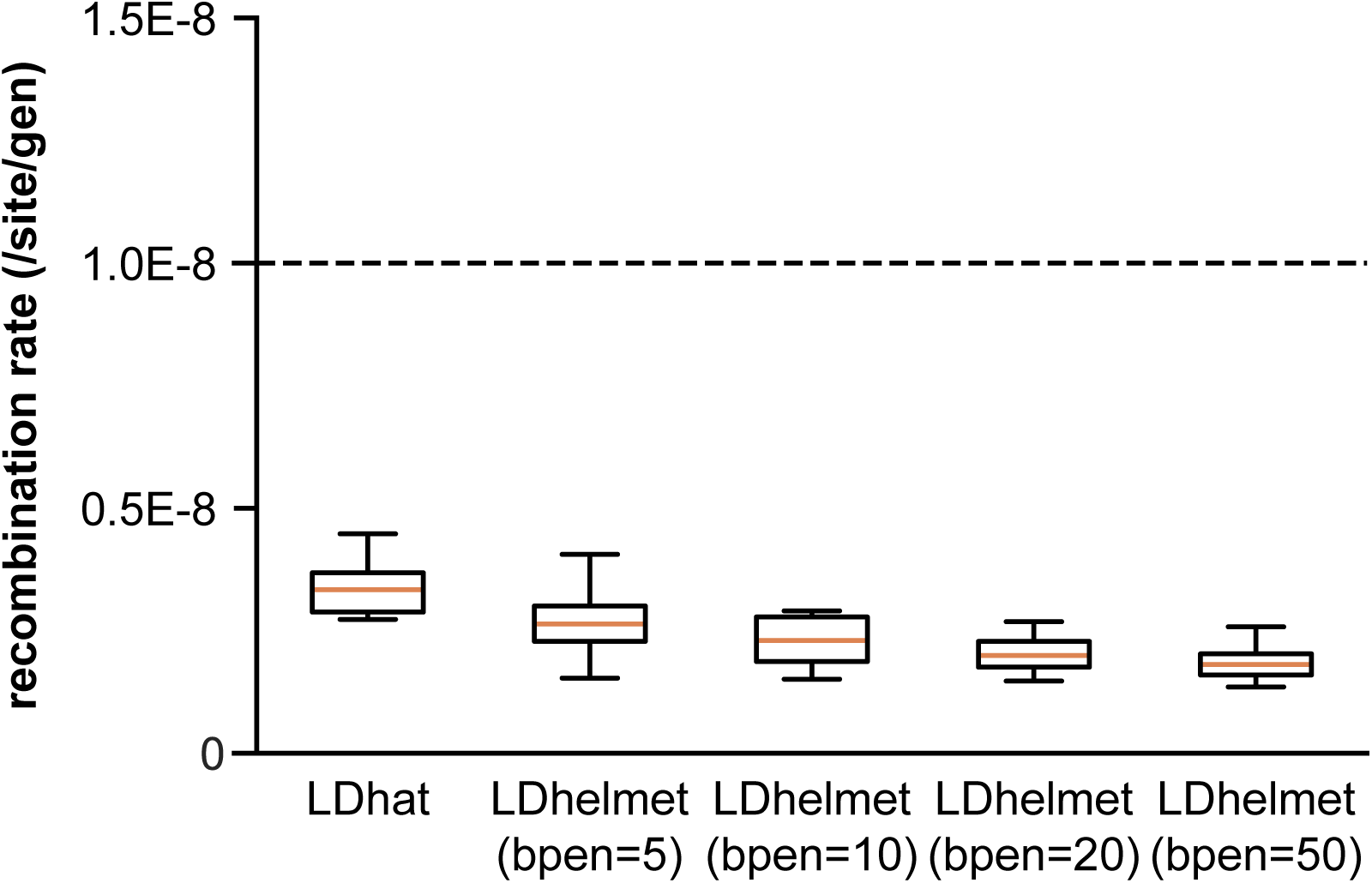
Performance of the recombination estimators. Performance of the recombination estimators LDhat and LDhelmet under the demographic history inferred in the coppery titi monkey by Terbot et al. (2026). The dashed line represents the constant recombination rate used in the simulations (10^-8^ /site/generation). Results for LDhelmet are shown for block penalties (bpen) of 5, 10, 20, and 50.

Having quantified the extent of expected mis-inference of recombination rates under the coppery titi monkey-specific population history, we then estimated the empirical landscapes of recombination. Given that LDhat and LDhelmet both infer the population-scaled recombination rate (ρ = 4𝑁_𝑒_𝑟), we calculated the effective population size, *N_e_*, from the empirically observed 𝜃 of 0.0043 in order to estimate the per-generation recombination rate, *r* (i.e., 𝑟 = ρ⁄4𝑁_𝑒_, with 𝑁_𝑒_ = 𝜃/4𝜇). Furthermore, due to the above-described expectation of under-estimation as well as uncertainties in *N_e_*, we re-scaled rates such that the total autosomal genetic map length was equal to that recently obtained from pedigreed individuals (Versoza et al. 2026b), whilst preserving the relative heterogeneity in recombination rates across the genome. Taking this approach, we inferred mean fine-scale genome wide recombination rates of 0.978 cM/Mb with LDhat (with sex-averaged rates ranging from 0.758 cM/Mb on one of the two longest autosomes, chromosome 12, to 1.177 cM/Mb on the shortest autosome, chromosome 22) and 0.975 cM/Mb with LDhelmet. Figure 3 provides the genome-wide recombination rates inferred by each method (and see Supplementary Figure S3 for the landscape of recombination across individual autosomes and Supplementary Figure S4 for the correlation between the two recombination maps).

**Figure 3.**
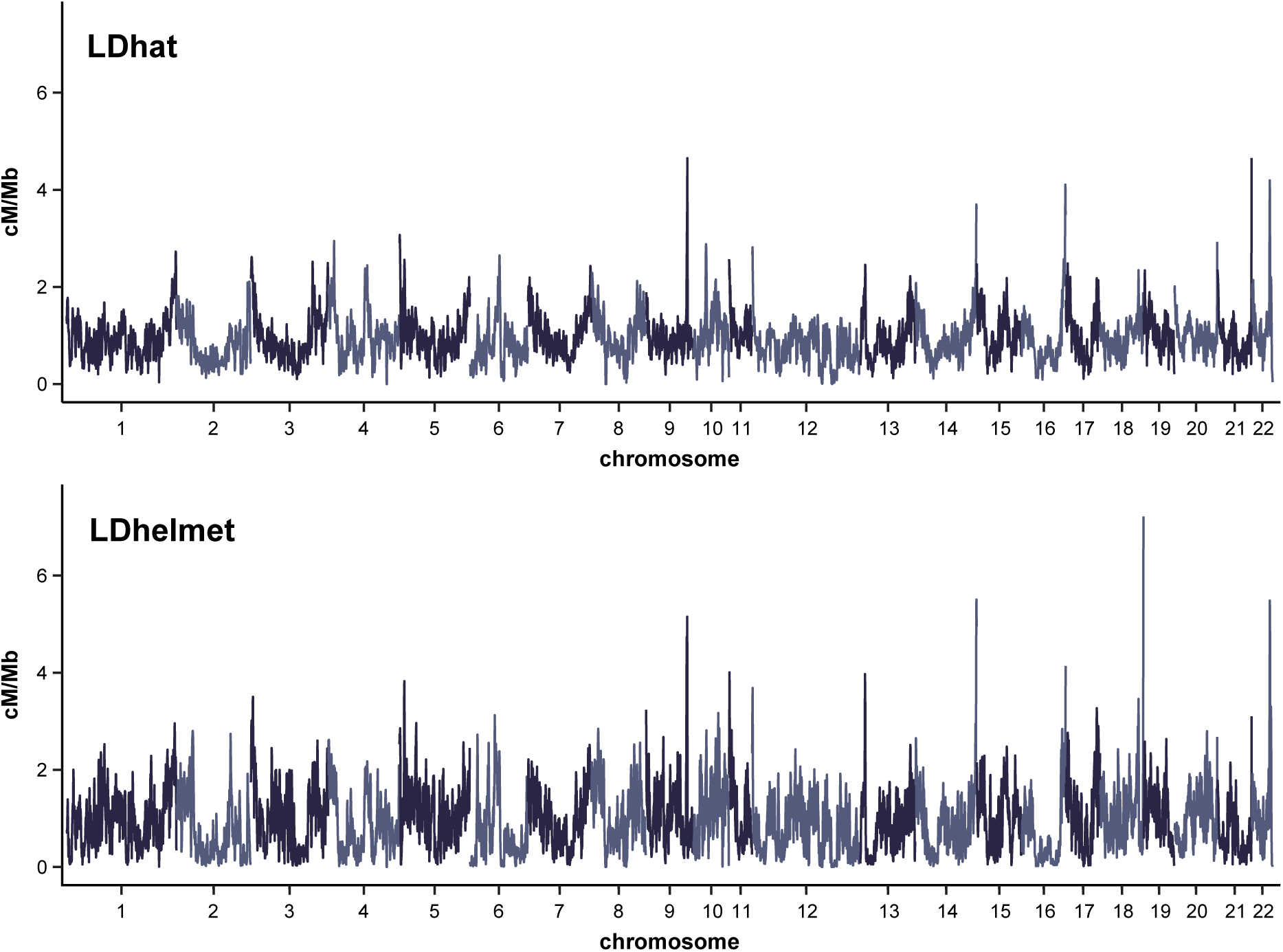
Fine-scale rates of recombination. Genome-wide recombination rates inferred using LDhat (top) and LDhelmet (bottom) for genomic windows of size 1 Mb, with a 500 kb step size (and see Supplementary Figure S3 for the heterogeneity in recombination rates across all autosomes). Results for LDhat and LDhelmet are shown for a block penalty of 5.

The genome-wide average rates inferred in coppery titi monkeys are thus similar to those inferred in another platyrrhine, common marmosets (0.91 cM/Mb; Soni, Versoza et al. 2025b), but higher than those previously inferred in several catarrhines of biomedical interest, including rhesus macaques (0.43 ± 0.33 cM/Mb; Xue et al. 2020) and vervet monkeys (0.43 ± 0.44 cM/Mb; Pfeifer 2020b). Moreover, these rates are within the same range as those previously reported in a number of great apes, including humans (1.32 ± 1.40 cM/Mb [International HapMap Consortium 2007], with an average rate of 0.945 cM/Mb in males and 1.518 cM/Mb in females [Halldorsson et al. 2019]), chimpanzees, bonobos, and gorillas (∼1.19 cM/Mb; Stevison et al. 2016). A note of caution is necessary, however, when comparing inferred recombination rates across different studies as the impact of demographic histories on these estimates have been considered to varying degrees between studies.

To further characterize the fine-scale distribution of recombination activity across the coppery titi monkey genome, we identified putative recombination hotspots using LDhot (Auton et al. 2014) and applied a multi-stage filtering procedure that integrates criteria from the Great Ape Recombination Project (Stevison et al. 2016) to construct a robust set of hotspot candidates. Following this filtering pipeline, we observed 9,210 hotspots, a number comparable to estimates previously obtained from non-human great apes (for which samples of similar size are available) using a modified LDhot framework (Nigerian chimpanzees: 9,316 hotspots; Western chimpanzees: 12,599 hotspots; gorillas: 10,384 hotspots; bonobos: 14,081 hotspots; see Table 3 in Stevison et al. 2016). Notably, the vast majority of hotspots (8,692, or 94.4%) contained the putative PRDM9 binding sequence recently identified through computational analyses (Versoza et al. 2026b) — a significant enrichment compared to the genomic background (background rate: 562 out of 9,210, or 6.1%; Fisher’s exact test: *p*-value ≍ 0).

Lastly, studying scale-specific covariation of recombination with a variety of genomic factors allowed us to investigate its impact on other evolutionary processes (Figure 4). In agreement with earlier studies, and concordant with one of the most prominent patterns in population genetics (Begun and Aquadro 1992), recombination in coppery titi monkeys is strongly positively correlated with nucleotide diversity, as expected from the effects of selection at linked sites reducing diversity in low recombination rate regions. A positive, albeit much weaker, correlation also exists with divergence, likely resulting from the mutagenic effects of recombination (Halldorsson et al. 2019). Nucleotide diversity and divergence are themselves strongly positively correlated due to the shared influences of genomic context (such as GC content and CpG density) and regional mutation rate variation (Hodgkinson and Eyre-Walker 2011). As expected from GC-biased gene conversion (Duret and Galtier 2009), recombination is weakly positively correlated with GC-density. In contrast, recombination rate and nucleotide diversity are both negatively correlated with gene density, likely driven by the preferential placement of PRDM9-dependent recombination within intergenic regions (Myers et al. 2005) as well as the pervasive effects of purifying and background selection (Charlesworth et al. 1993). A negative correlation with repeat content was also observed, consistent with the accumulation of repetitive elements in heterochromatic regions where recombination is suppressed, likely reflecting structural and epigenetic constraints that promote genome stability (see the review by Charlesworth et al. 1994). Notably, nearly all correlations exhibit a pronounced dependence on genomic scale, with the correlations observed at fine scales being consistent with the transient and rapidly evolving nature of PRDM9-dependent recombination hotspots, and the correlations observed at the broad scales reflecting the cumulative effects of hotspot turnover, genome organization, and long-term constraints on recombination placement. The emergence of stronger recombination–diversity and recombination–divergence correlations at the broad-scale therefore suggests that, while PRDM9 determines the fine-scale localization of recombination events, the evolutionary consequences of recombination primarily manifest at broader genomic scales — in agreement with both theoretical expectations and empirical observations in other primates (e.g., Auton et al. 2012; Pfeifer and Jensen 2016; Stevison et al. 2016; Pfeifer 2020b; and see the review of Cutter and Payseur 2013).

**Figure 4.**
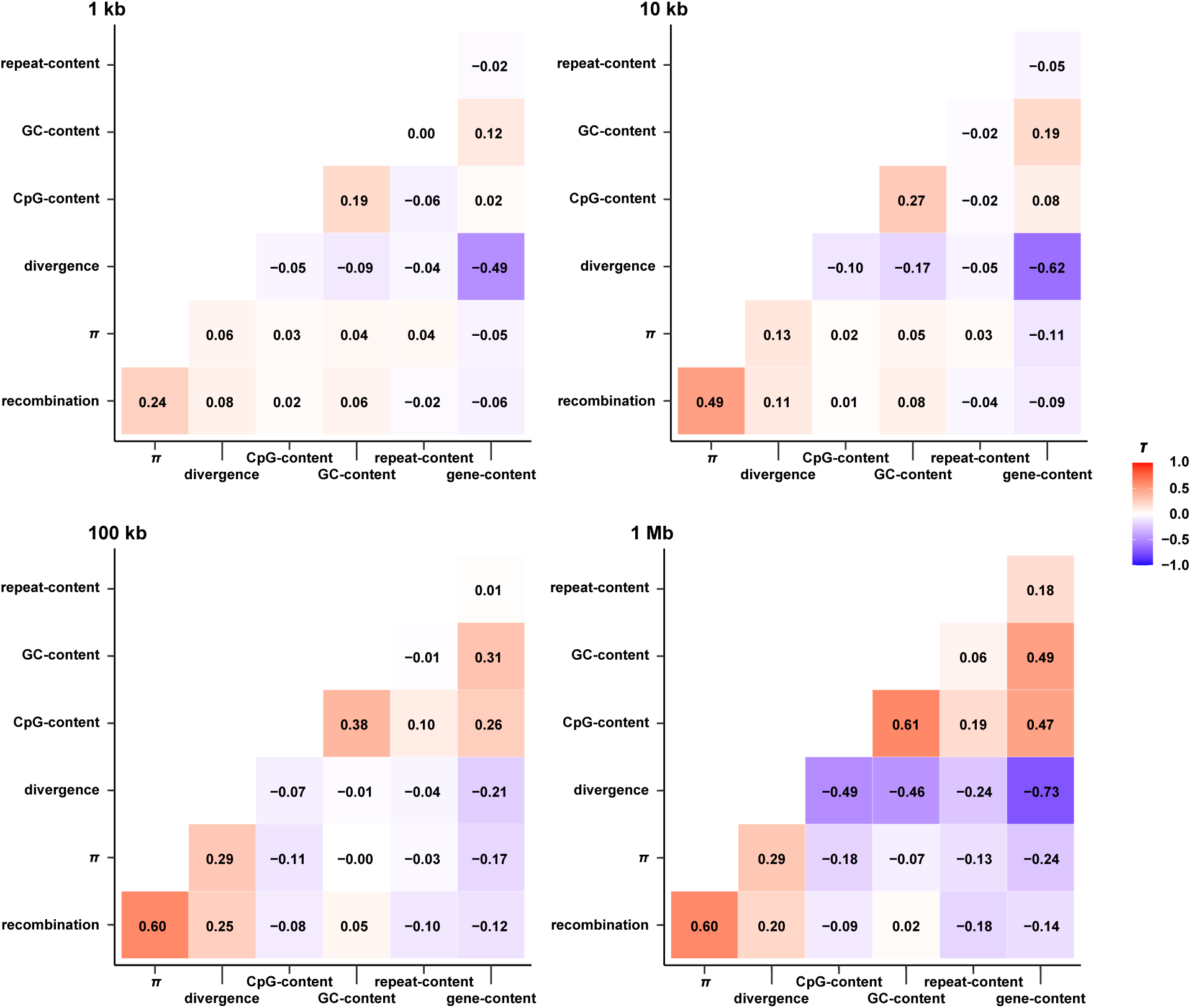
Correlation between recombination rates and genomic features. Correlation between recombination rates and genomic features — namely nucleotide diversity (*π*), neutral divergence, CpG-content, GC-content, repeat-content, and gene-content — calculated across a variety of window sizes (1 kb, 10 kb, 100 kb, and 1 Mb). Partial Kendall’s τ correlations are color-coded, with a red coloring indicating positive correlations and a blue coloring indicating negative correlations. Color intensity is proportional to the strength of correlation.

## CONCLUDING THOUGHTS

This study presents the first fine-scale, genome-wide mutation and recombination rate maps in the coppery titi monkey, *P. cupreus*. Interestingly, rates were generally more consistent with estimates in the great apes, rather than (the admittedly sparse) estimates previously reported from other platyrrhines. This work thus again highlights the important levels of rate heterogeneity even amongst relatively closely related species, and highlights the need for more dense species- and population-sampling across the primate clade. Given the importance of coppery titi monkeys as a model system of neurobiology and social behavior, these estimated rate landscapes will prove useful for future research, including for example when performing genomic scans for selection, as well as for interpreting genome-wide association studies, for traits of biomedical interest.

## ACKNOWLEDGEMENTS

Computations were performed on the Sol supercomputer at Arizona State University (Jennewein et al. 2023).

## REFERENCES

1000 Genomes Project Consortium. 2015. A global reference for human genetic variation. Nature. 526(7571):68–74.

Armstrong J, Hickey G, Diekhans M, Fiddes IT, Novak AM, Deran A, Fang Q, Xie D, Feng S, Stiller J, et al. 2020. Progressive Cactus is a multiple-genome aligner for the thousand-genome era. Nature. 587(7833):246–251.

Auton A, Fledel-Alon A, Pfeifer S, Venn O, Ségurel L, Street T, Leffler EM, Bowden R, Aneas I, Broxholme J, et al. 2012. A fine-scale chimpanzee genetic map from population sequencing. Science. 336(6078):193–198.

Auton A, McVean G. 2007. Recombination rate estimation in the presence of hotspots. Genome Res.17(8):1219–1227.

Auton A, Myers S, McVean G. 2014. Identifying recombination hotspots using population genetic data. arXiv [Preprint] 10.48550/arXiv.1403.4264

Baer CF, Miyamoto MM, Denver DR. 2007. Mutation rate variation in multicellular eukaryotes: causes and consequences. Nat Rev Genet. 8(8):619–631.

Bales KL, Ardekani CS, Baxter A, Karaskiewicz CL, Kusek JX, Lau AR, Savidge LE, Sayler, KR, Witczak LR. 2021. What is a pair bond? Horm Behav. 136:105062.

Bales KL, Mason WA, Catana C, Cherry SR, Mendoza SP. 2007. Neural correlates of pair-bonding in a monogamous primate. Brain Res. 1184(1):245–253.

Baumdicker F, Bisschop G, Goldstein D, Gower G, Ragsdale AP, Tsambos G, Zhu S, Eldon B, Ellerman EC, Galloway JG, et al. 2022. Efficient ancestry and mutation simulation with msprime 1.0. Genetics. 220(3):iyab229.

Begun DJ, Aquadro CF. 1992. Levels of naturally occurring DNA polymorphism correlate with recombination rates in *D. melanogaster*. Nature. 356(6369):519–520.

Bergeron LA, Besenbacher S, Turner TN, Versoza CJ, Wang RJ, Price AL, Armstrong E, Riera M, Carlson J, Chen HY, et al. 2022. The Mutationathon highlights the importance of reaching standardization in estimates of pedigree-based germline mutation rates. eLife. 11:e73577.

Besenbacher S, Hvilsom C, Marques-Bonet T, Mailund T, Schierup MH. 2019. Direct estimation of mutations in great apes reconciles phylogenetic dating. Nat Ecol Evol. 3(2):286–292.

Brazier T, Glémin S. 2024. Diversity in recombination hotspot characteristics and gene structure shape fine-scale recombination patterns in plant genomes. Mol Biol Evol. 41(9):msae183.

Campbell CR, Tiley GP, Poelstra JW, Hunnicutt KE, Larsen PA, Lee HJ, Thorne JL, Dos Reis M, Yoder AD. 2021. Pedigree-based and phylogenetic methods support surprising patterns of mutation rate and spectrum in the gray mouse lemur. Heredity (Edinb). 127(2):233–244.

Chan AH, Jenkins PA, Song YS. 2012. Genome-wide fine-scale recombination rate variation in *Drosophila melanogaster*. PLoS Genet. 8(12):e1003090.

Charlesworth B, Jensen JD. 2021. Effects of selection at linked sites on patterns of genetic variability. Annu Rev Ecol Evol Syst. 52:177–197.

Charlesworth B, Jensen JD. 2022. How can we resolve Lewontin’s Paradox? Genome Biol Evol. 14(7):evac096.

Charlesworth B, Morgan MT, Charlesworth D. 1993. The effect of deleterious mutations on neutral molecular variation. Genetics. 134(4):1289–1303.

Charlesworth B, Sniegowski P, Stephan W. 1994. The evolutionary dynamics of repetitive DNA in eukaryotes. Nature. 371(6494):215–220.

Chen S, Zhou Y, Chen Y, Gu J. 2018. fastp: an ultra-fast all-in-one FASTQ preprocessor. Bioinformatics. 34(17):i884–i890.

Chintalapati M, Moorjani P. 2020. Evolution of the mutation rate across primates. Curr Opin in Genet Dev. 62:58–64.

Clark AG, Wang X, Matise T. 2010. Contrasting methods of quantifying fine structure of human recombination. Annu Rev Genomics Hum Genet. 11:45–64.

Conley AJ, Berger T, Del Razo RA, Cotterman RF, Sahagún E, Goetze LR, Jacob S, Weinstein TAR, Dufek ME, Mendoza SP, et al. 2022. The onset of puberty in colony-housed male and female titi monkeys (*Plecturocebus cupreus*): possible effects of oxytocin treatment during peri-adolescent development. Horm Behav. 142:105157.

Cutter AD, Payseur BA. 2013. Genomic signatures of selection at linked sites: unifying the disparity among species. Nat Rev Genet. 14(4):262–274.

Danecek P, Bonfield JK, Liddle J, Marshall J, Ohan V, Pollard MO, Whitwham A, Keane T, McCarthy SA, Davies RM, et al. 2021. Twelve years of SAMtools and BCFtools. Giga Science. 10(2):giab008.

Dapper AL, Payseur BA. 2018. Effects of demographic history on the detection of recombination hotspots from linkage disequilibrium. Mol Biol Evol. 35(2):335–353.

de Magalhães JP, Costa J. 2009. A database of vertebrate longevity records and their relation to other life-history traits. J Evol Biol. 22(8):1770–1774.

Duret L, Galtier N. 2009. Biased gene conversion and the evolution of mammalian genomic landscapes. Annu Rev Genomics Hum Genet. 285–311.

Dutheil JY. 2024. On the estimation of genome-average recombination rates. Genetics. 227(2):iyae051.

Eggertsson HP, Jonsson H, Kristmundsdottir S, Hjartarson E, Kehr B, Masson G, Zink F, Hjorleifsson KE, Jonasdottir A, Jonasdottir A, et al. 2017. Graphtyper enables population-scale genotyping using pangenome graphs. Nat Genet. 49(11):1654– 1660.

Felsenstein J. 1974. The evolutionary advantage of recombination. Genetics. 78(2):737–756.

Ghafoor S, Santos J, Versoza CJ, Jensen JD, Pfeifer SP. 2023. The impact of sample size and population history on observed mutational spectra: a case study in human and chimpanzee populations. Genome Biol Evol. 15(3):evad019.

Glazko GV, Nei M. 2003. Estimation of divergence times for major lineages of primate species. Mol Biol Evol. 20(3):424–434.

Grant CE, Bailey TL, Noble WS. 2011. FIMO: scanning for occurrences of a given motif. Bioinformatics. 27(7):1017–1018.

Haldane JBS. 1932. The causes of evolution. Longmans, Green, & Co, London, UK.

Haldane JBS. 1935. The rate of spontaneous mutation of a human gene. J Genet. 31:317–326.

Halldorsson BV, Palsson G, Stefansson OA, Jonsson H, Hardarson MT, Eggertsson HP, Gunnarsson B, Oddsson A, Halldorsson GH, Zink F, et al. 2019. Characterizing mutagenic effects of recombination through a sequence-level genetic map. Science. 363(6425):eaau1043.

Hickey G, Paten B, Earl D, Zerbino D, Haussler D. 2013. HAL: a hierarchical format for storing and analyzing multiple genome alignments. Bioinformatics. 29(10):1341–1342.

Hill WG, Robertson A. 1966. The effect of linkage on limits to artificial selection. Genet Res. 8(3):269–294.

Hodgkinson A, Eyre-Walker A. 2011. Variation in the mutation rate across mammalian genomes. Nat Rev Genet. 12(11):756–766.

International HapMap Consortium. 2007. A second generation human haplotype map of over 3.1 million SNPs. Nature. 449(7164):851–861.

Jennewein DM, Lee J, Kurtz C, Dizon W, Shaeffer I, Chapman A, Chiquete A, Burks J, Carlson A, Mason N, et al. 2023. The Sol supercomputer at Arizona State University. In Practice and Experience in Advanced Research Computing 2023: Computing for the Common Good (PEARC ’23). Association for Computing Machinery, New York, NY, USA, 296–301.

Johnston S. 2024. Understanding the genetic basis of variation in meiotic recombination: past, present, and future. Mol Biol Evol. 41(7):msae112.

Johri P, Aquadro CF, Beaumont M, Charlesworth B, Excoffier L, Eyre-Walker A, Keightley PD, Lynch M, McVean G, Payseur BA, et al. 2022. Recommendations for improving statistical inference in population genomics. PLoS Biol. 20(5):e3001669.

Johri P, Charlesworth B, Jensen JD. 2020. Toward an evolutionarily appropriate null model: jointly inferring demography and purifying selection. Genetics. 215(1):173– 192.

Jónsson H, Sulem P, Kehr B, Kristmundsdottir S, Zink F, Hjartarson E, Hardarson MT, Hjorleifsson KE, Eggertsson HP, Gudjonsson SA, et al. 2017. Parental influence on human germline *de novo* mutations in 1,548 trios from Iceland. Nature. 549(7673):519–522.

Kimura M. 1968. Evolutionary rate at the molecular level. Nature. 217(5129):624– 626.

Kimura M. 1983. The Neutral Theory of Molecular Evolution. 1st ed. Cambridge University Press. Available from: https://www.cambridge.org/core/product/identifier/9780511623486/type/book

Kong A, Gudbjartsson DF, Sainz J, Jonsdottir GM, Gudjonsson SA, Richardsson B, Sigurdardottir S, Barnard J, Hallbeck B, Masson G, et al. 2002. A high-resolution recombination map of the human genome. Nat Genet. 31(3):241–247.

Kuderna LFK, Ulirsch JC, Rashid S, Ameen M, Sundaram L, Hickey G, Cox AJ, Gao H, Kumar A, Aguet F, et al. 2024. Identification of constrained sequence elements across 239 primate genomes. Nature. 625(7996):735–742.

Lau AR, Baxter A, He S, Loyant L, Ortiz-Jimenez CA, Bauman MD, Bales KL, Freeman SM. 2024. Age, pair tenure and parenting, but not face identity, predict looking behaviour in a pair-bonded South American primate. Anim Behav. 217:53– 63.

Li H. 2013. Aligning sequence reads, clone sequences and assembly contigs with BWA-MEM. arXiv [Preprint] 10.48550/arXiv.1303.3997

Lynch M. 2010. Evolution of the mutation rate. Trends Genet. 26(8):345–352.

Martin M, Ebert P, Marschall T. 2023. Read-based phasing and analysis of phased variants with WhatsHap. Methods Mol Biol. 2590:127–138.

Maynard Smith J, Haigh J. 1974. The hitch-hiking effect of a favourable gene. Genet Res. 23(1):23–35.

McVean G, Awadalla P, Fearnhead P. 2002. A coalescent-based method for detecting and estimating recombination from gene sequences. Genetics. 160(3):1231–1241.

McVean GAT, Myers SR, Hunt S, Deloukas P, Bentley DR, Donnelly P. 2004. The fine-scale structure of recombination rate variation in the human genome. Science. 304(5670):581–584.

Myers S, Bottolo L, Freeman C, McVean G, Donnelly P. 2005. A fine-scale map of recombination rates and hotspots across the human genome. Science. 310(5746):321–324.

Pacifici M, Santini L, Di Marco M, Baisero D, Francucci L, Grottolo Marasini G, Visconti P, Rondinini C. 2013. Generation length for mammals. Nat Conserv. 5:87– 94.

Peñalba JV, Wolf JBW. 2020. From molecules to populations: appreciating and estimating recombination rate variation. Nat Rev Genet. 21(8):476–492.

Perez SI, Tejedor MF, Novo NM, Aristide L. 2013. Divergence times and the evolutionary radiation of New World monkeys (Platyrrhini, primates): an analysis of fossil and molecular data. PLoS One. 8(6):e68029.

Pfeifer SP. 2017. From next-generation resequencing reads to a high-quality variant data set. Heredity (Edinb). 118(2):111–124.

Pfeifer SP. 2020a. Spontaneous mutation rates. In Ho SYW (ed). The Molecular Evolutionary Clock. Theory and Practice. Springer Nature, pp. 35–44.

Pfeifer SP. 2020b. A fine-scale genetic map for vervet monkeys. Mol Biol Evol. 37(7):1855–1865.

Pfeifer SP. 2021. Studying mutation rate evolution in primates-the effects of computational pipelines and parameter choices. Giga Science. 10(10):giab069.

Pfeifer SP, Baxter A, Savidge LE, Sedlazeck FJ, Bales KL. 2024. *De novo* genome assembly for the coppery titi monkey (*Plecturocebus cupreus*): an emerging nonhuman primate model for behavioral research. Genome Biol Evol. 16(5):evae108.

Pfeifer SP, Jensen JD. 2016. The impact of linked selection in chimpanzees: a comparative study. Genome Biol Evol. 8(10):3202–3208.

Quinlan AR, Hall IM. 2010. BEDTools: a flexible suite of utilities for comparing genomic features. Bioinformatics 26(6):841–842.

Ritz RR, Noor MAF, Singh ND. 2017. Variation in recombination rate: adaptive or not? Trends Genet. 33(5):364–374.

Samuk K, Noor MAF. 2022. Gene flow biases population genetic inference of recombination rate. G3 (Bethesda). 12(11):jkac236.

Soni V, Jensen JD. 2024. Temporal challenges in detecting balancing selection from population genomic data. G3 (Bethesda). 14(6):jkae069.

Soni V, Johri P, Jensen JD. 2023. Evaluating power to detect recurrent selective sweeps under increasingly realistic evolutionary null models. Evolution. 77(10):2113–2127.

Soni V, Pfeifer SP, Jensen JD. 2024. The effect of mutation and recombination rate heterogeneity on the inference of demography and the distribution of fitness effects. Genome Biol Evol. 16(2):evae004.

Soni V, Pfeifer SP, Jensen JD. 2025. Recent insights into the evolutionary genomics of the critically endangered aye-aye (*Daubentonia madagascariensis*). Am J Primatol. 87(12):e70105.

Soni V, Versoza CJ, Jensen JD, Pfeifer SP. 2025b. Inferring the landscapes of mutation and recombination in the common marmoset (*Callithrix jacchus*) in the presence of twinning and hematopoietic chimerism. BioRxiv [Preprint] 10.1101/2025.07.01.662565

Soni V, Versoza CJ, Terbot JW, Jensen JD, Pfeifer SP. 2025a. Inferring fine-scale mutation and recombination rate maps in aye-ayes (*Daubentonia madagascariensis*). Ecol Evol. 15(11):e72314.

Stapley J, Feulner PGD, Johnston SE, Santure AW, Smadja CM. 2017. Variation in recombination frequency and distribution across eukaryotes: patterns and processes. Philos Trans R Soc Lond B Biol Sci. 372(1736):20160455.

Stevison LS, Woerner AE, Kidd JM, Kelley JL, Veeramah KR, McManus KF, Great Ape Genome Project, Bustamante CD, Hammer MF, Wall JD. 2016. The time scale of recombination rate evolution in great apes. Mol Biol Evol. 33(4):928–945.

Stumpf MPH, McVean GAT. 2003. Estimating recombination rates from population-genetic data. Nat Rev Genet. 4(12):959–968.

Tatsumoto S, Go Y, Fukuta K, Noguchi H, Hayakawa T, Tomonaga M, Hirai H, Matsuzawa T, Agata K, Fujiyama A. 2017. Direct estimation of *de novo* mutation rates in a chimpanzee parent-offspring trio by ultra-deep whole genome sequencing. Sci Rep. 7(1):13561.

Terbot JW, Soni V, Versoza CJ, Bales KL, Pfeifer SP, Jensen JD. 2026. Inferring the demographic history of coppery titi monkeys (*Plecturocebus cupreus*) from high-quality, whole-genome, population-level data. BioRxiv [Preprint] 10.64898/2026.01.09.698678.

Terbot JW, Soni V, Versoza CJ, Milhaven M, Calahorra-Oliart A, Shah D, Pfeifer SP, Jensen JD. 2025. Interpreting patterns of X chromosomal relative to autosomal diversity in ayes-ayes (*Daubentonia madagascariensis*). Am J Primatol. 87(12):e70091.

Tran LA, Pfeifer SP. 2018. Germ line mutation rates in Old World monkeys. In: John Wiley & Sons, Ltd, editor. eLS. 1st ed. Wiley. pp. 1–10.

Valeggia CR, Mendoza SP, Fernandez-Duque E, Mason WA, Lasley B. 1999. Reproductive biology of female titi monkeys (*Callicebus moloch*) in captivity. Am J Primatol. 47(3):183–195.

Van Belle S, Fernandez-Duque E, Di Fiore A. 2016. Demography and life history of wild red titi monkeys (*Callicebus discolor*) and equatorial sakis (*Pithecia aequatorialis*) in Amazonian Ecuador: a 12-year study. Am J Primatol. 78(2):204– 215.

van der Auwera GA, O’Connor BD. 2020. Genomics in the cloud: using Docker, GATK, and WDL in Terra. Sebastopol: O’Reilly Media.

Venn O, Turner I, Mathieson I, de Groot N, Bontrop R, McVean G. 2014. Strong male bias drives germline mutation in chimpanzees. Science. 344(6189):1272–1275.

Versoza CJ, Bales KL, Jensen JD, Pfeifer SP. 2026a. Characterizing the rates and patterns of *de novo* germline mutations in coppery titi monkeys (*Plecturocebus cupreus*). BioRxiv [Preprint]

Versoza CJ, Bales KL, Jensen JD, Pfeifer SP. 2026b. Sex-specific landscapes of crossover and non-crossover recombination in coppery titi monkeys (*Plecturocebus cupreus*). BioRxiv [Preprint] 10.64898/2026.01.11.698870.

Versoza CJ, Ehmke EE, Jensen JD, Pfeifer SP. 2025. Characterizing the rates and patterns of *de novo* germline mutations in the aye-aye (*Daubentonia madagascariensis*). Mol Biol Evol. 42(3):msaf034.

Versoza CJ, Lloret-Villas A, Jensen JD, Pfeifer SP. 2025. A pedigree-based map of crossovers and noncrossovers in aye-ayes (*Daubentonia madagascariensis*). Genome Biol Evol. 17(5):evaf072.

Versoza C, Weiss S, Johal R, La Rosa B, Jensen JD, Pfeifer SP. 2024. Novel insights into the landscape of crossover and non-crossover events in rhesus macaques (*Macaca mulatta*). Genome Biol Evol. 16(1):evad223.

Virtanen P, Gommers R, Oliphant TE, Haberland M, Reddy T, Cournapeau D, Burovski E, Peterson P, Weckesser W, Bright J, et al. 2020. SciPy 1.0: fundamental algorithms for scientific computing in Python. Nat Methods. 17(3):261–272.

Wall JD, Robinson JA, Cox LA. 2022. High-resolution estimates of crossover and noncrossover recombination from a captive baboon colony. Genome Biol Evol. 14(4):evac040.

Xue C, Rustagi N, Liu X, Raveendran M, Harris RA, Venkata MG, Rogers J, Yu F. 2020. Reduced meiotic recombination in rhesus macaques and the origin of the human recombination landscape. PLoS One. 15(8):e0236285.

Zablocki-Thomas P, Rebout N, Karaskiewicz CL, Bales KL. 2023. Survival rates and mortality risks of *Plecturocebus cupreus* at the California National Primate Research Center. Am J Primatol. 85(10):e23531.

Zoonomia Consortium. 2020. A comparative genomics multitool for scientific discovery and conservation. Nature. 587(7833):240–245.

