## Supplementary Materials for "Inferring fine-scale rates of mutation and recombination in the coppery titi monkey (*Plecturocebus cupreus*)"

| chromosome | length | # SNPs | Ts/Tv | # invariable sites |
|---|---|---|---|---|
| 1 | 235,103,535 | 644,311 | 2.64 | 148,205,777 |
| 2 | 158,738,102 | 434,111 | 2.70 | 98,497,389 |
| 3 | 165,937,890 | 467,368 | 2.56 | 102,000,154 |
| 4 | 151,623,469 | 383,446 | 2.65 | 94,167,465 |
| 5 | 150,993,727 | 430,053 | 2.60 | 94,005,171 |
| 6 | 123,740,199 | 345,869 | 2.69 | 76,193,309 |
| 7 | 135,512,294 | 367,789 | 2.52 | 82,843,393 |
| 8 | 116,834,941 | 346,007 | 2.47 | 72,431,464 |
| 9 | 101,234,152 | 273,083 | 2.58 | 60,149,663 |
| 10 | 75,878,286 | 213,058 | 2.77 | 44,393,010 |
| 11 | 49,373,254 | 131,462 | 2.66 | 29,936,619 |
| 12 | 231,176,762 | 586,920 | 2.66 | 141,745,863 |
| 13 | 118,029,634 | 282,023 | 2.80 | 73,673,212 |
| 14 | 130,776,705 | 283,266 | 2.71 | 81,241,012 |
| 15 | 95,533,141 | 249,431 | 2.58 | 59,771,475 |
| 16 | 94,172,006 | 247,341 | 2.59 | 59,156,815 |
| 17 | 73,180,778 | 194,592 | 2.83 | 44,439,300 |
| 18 | 92,880,398 | 286,221 | 2.53 | 57,538,597 |
| 19 | 65,768,754 | 198,900 | 2.91 | 42,772,996 |
| 20 | 91,752,637 | 241,048 | 2.42 | 54,790,595 |
| 21 | 73,432,007 | 182,368 | 2.55 | 44,848,075 |
| 22 | 44,717,611 | 116,987 | 2.52 | 27,199,013 |
| $\sum$ or $\varnothing$ | **2,576,390,282** | **6,905,654** | **2.63** | **1,590,000,367** |

**Supplementary Table S1.** Population genomic data.

| | accessibility threshold | | | mutation rates | | | | | |
|---|---|---|---|---|---|---|---|---|---|
| window size | min. accessible length (bp) | number of windows that meet threshold | proportion of windows that meet threshold | 6 year generation time | | | 9 year generation time | | |
| | | | | 32 mya | 33 mya | 36 mya | 32 mya | 33 mya | 36 mya |
| 1 Mb | 10,000 | 4,295 | 0.857 | 1.051E-08 | 1.019E-08 | 9.344E-09 | 1.577E-08 | 1.529E-08 | 1.402E-08 |
| | 25,000 | 4,163 | 0.830 | 1.049E-08 | 1.017E-08 | 9.327E-09 | 1.574E-08 | 1.526E-08 | 1.399E-08 |
| | 50,000 | 3,975 | 0.793 | 1.048E-08 | 1.016E-08 | 9.314E-09 | 1.572E-08 | 1.524E-08 | 1.397E-08 |
| | 75,000 | 3,741 | 0.746 | 1.049E-08 | 1.017E-08 | 9.322E-09 | 1.573E-08 | 1.525E-08 | 1.398E-08 |
| | 100,000 | 3,428 | 0.684 | 1.050E-08 | 1.018E-08 | 9.336E-09 | 1.575E-08 | 1.528E-08 | 1.400E-08 |
| | 250,000 | 1,589 | 0.317 | 1.064E-08 | 1.032E-08 | 9.461E-09 | 1.597E-08 | 1.548E-08 | 1.419E-08 |
| 100 kb | 1,000 | 37,164 | 0.923 | 1.045E-08 | 1.013E-08 | 9.290E-09 | 1.568E-08 | 1.520E-08 | 1.393E-08 |
| | 2,500 | 35,493 | 0.882 | 1.044E-08 | 1.012E-08 | 9.280E-09 | 1.566E-08 | 1.518E-08 | 1.392E-08 |
| | 5,000 | 33,155 | 0.824 | 1.045E-08 | 1.014E-08 | 9.292E-09 | 1.568E-08 | 1.520E-08 | 1.394E-08 |
| | 7,500 | 30,968 | 0.769 | 1.048E-08 | 1.016E-08 | 9.315E-09 | 1.572E-08 | 1.524E-08 | 1.397E-08 |
| | 10,000 | 28,894 | 0.718 | 1.051E-08 | 1.019E-08 | 9.339E-09 | 1.576E-08 | 1.528E-08 | 1.401E-08 |
| | 25,000 | 17,274 | 0.429 | 1.063E-08 | 1.031E-08 | 9.452E-09 | 1.595E-08 | 1.547E-08 | 1.418E-08 |
| 10 kb | 100 | 254,635 | 0.977 | 1.052E-08 | 1.020E-08 | 9.351E-09 | 1.578E-08 | 1.530E-08 | 1.403E-08 |
| | 250 | 249,129 | 0.956 | 1.051E-08 | 1.020E-08 | 9.346E-09 | 1.577E-08 | 1.529E-08 | 1.402E-08 |
| | 500 | 241,325 | 0.926 | 1.051E-08 | 1.019E-08 | 9.343E-09 | 1.577E-08 | 1.529E-08 | 1.401E-08 |
| | 750 | 234,081 | 0.898 | 1.051E-08 | 1.020E-08 | 9.346E-09 | 1.577E-08 | 1.529E-08 | 1.402E-08 |
| | 1,000 | 226,709 | 0.870 | 1.052E-08 | 1.020E-08 | 9.349E-09 | 1.578E-08 | 1.530E-08 | 1.402E-08 |
| | 2,500 | 174,243 | 0.668 | 1.056E-08 | 1.024E-08 | 9.384E-09 | 1.583E-08 | 1.535E-08 | 1.408E-08 |
| 1 kb | 10 | 1,933,294 | 0.971 | 1.074E-08 | 1.042E-08 | 9.548E-09 | 1.611E-08 | 1.562E-08 | 1.432E-08 |
| | 25 | 1,890,310 | 0.949 | 1.068E-08 | 1.035E-08 | 9.490E-09 | 1.602E-08 | 1.553E-08 | 1.424E-08 |
| | 50 | 1,838,381 | 0.923 | 1.063E-08 | 1.031E-08 | 9.450E-09 | 1.595E-08 | 1.546E-08 | 1.417E-08 |
| | 75 | 1,792,451 | 0.900 | 1.061E-08 | 1.028E-08 | 9.428E-09 | 1.591E-08 | 1.543E-08 | 1.414E-08 |
| | 100 | 1,748,751 | 0.878 | 1.059E-08 | 1.027E-08 | 9.414E-09 | 1.589E-08 | 1.541E-08 | 1.412E-08 |
| | 250 | 1,470,597 | 0.738 | 1.053E-08 | 1.022E-08 | 9.364E-09 | 1.580E-08 | 1.532E-08 | 1.405E-08 |

**Supplementary Table S2.** Mean mutation rates for a variety of window sizes, accessibility thresholds, generation times, and *P. cupreus–H. sapiens* divergence times.

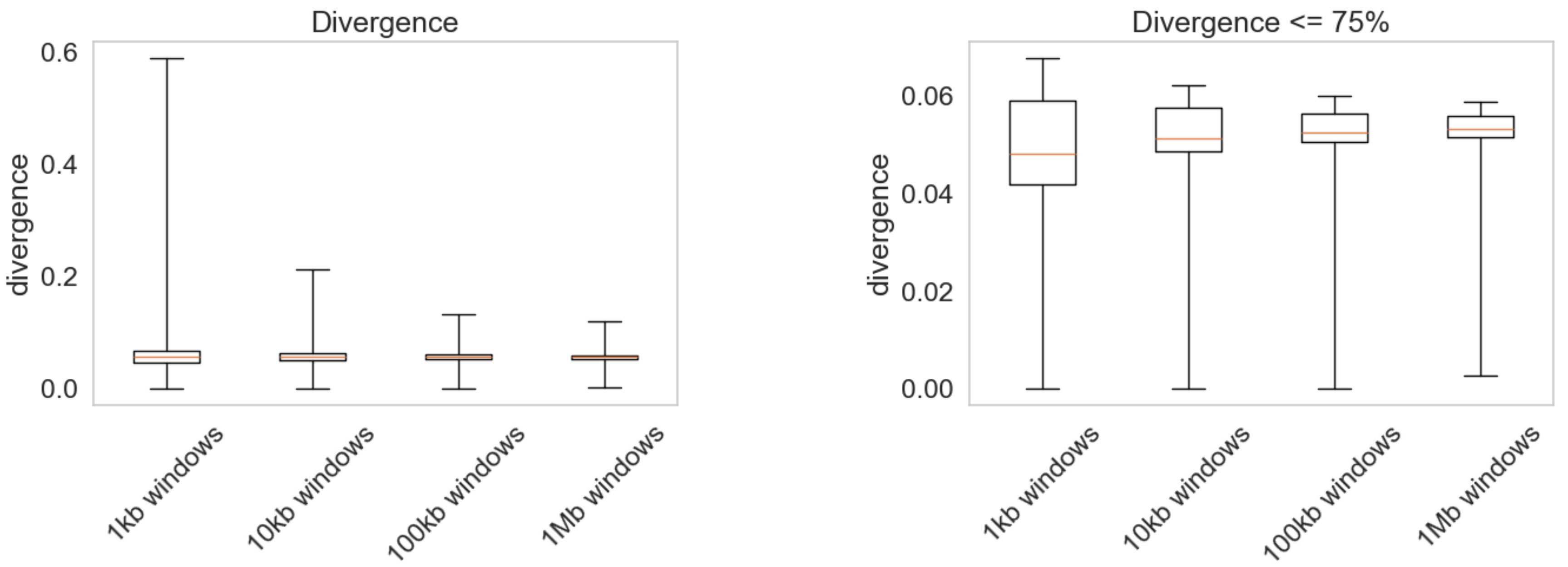


**Supplementary Figure S1.** Distributions of neutral divergence between *P. cupreus* and *H. sapiens* for all 1 kb, 10 kb, 100 kb and 1 Mb genomic windows (left) and for windows in the lower three quartiles (right). Orange bars represent the median neutral divergence, boxes represent the first and third quartiles, and whiskers represent the minimum and maximum values.

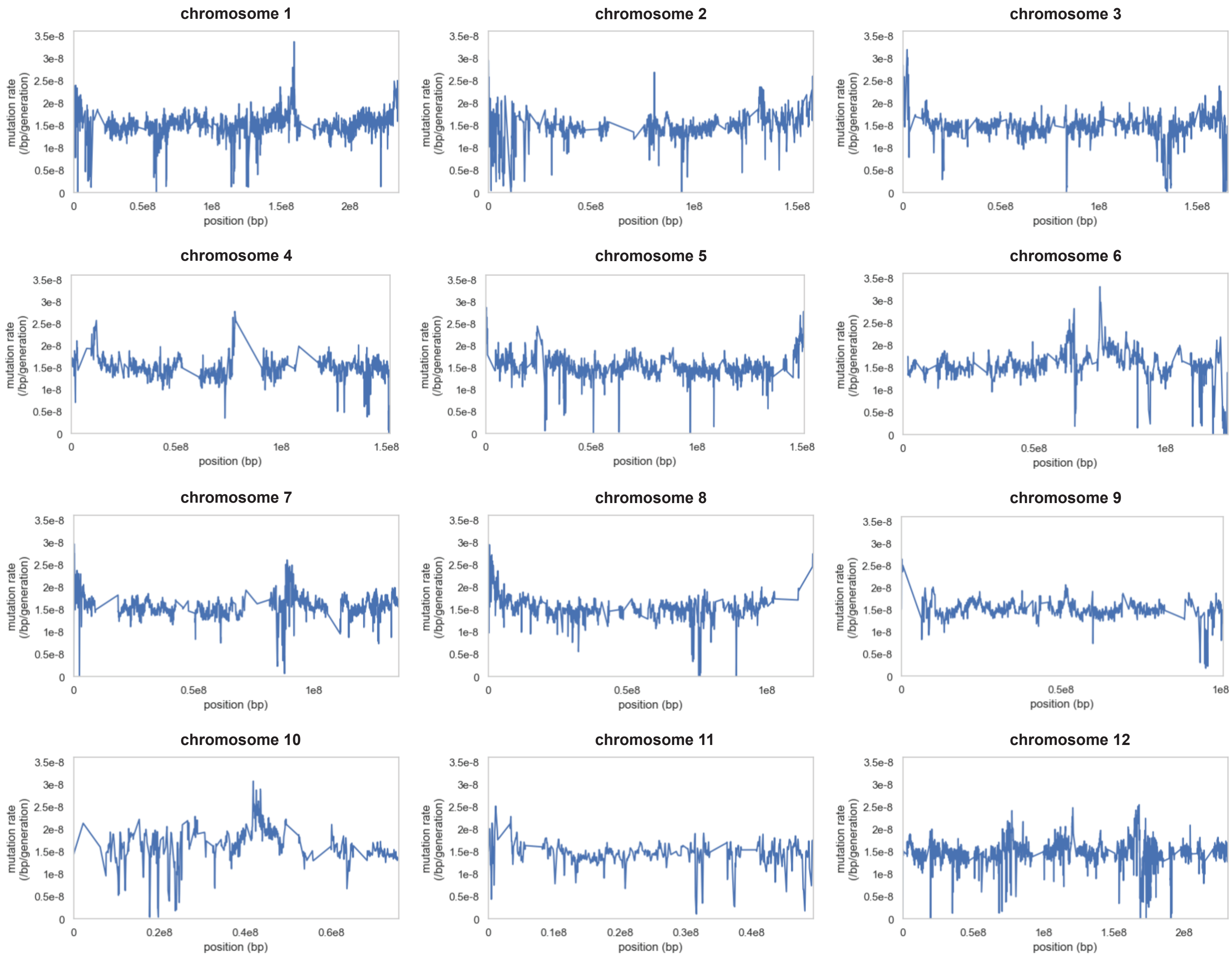
chromosome 1
chromosome 2
chromosome 3
chromosome 4
chromosome 5
chromosome 6
chromosome 7
chromosome 8
chromosome 9
chromosome 10
chromosome 11
chromosome 12
mutation rate (/bp/generation)
position (bp)
0
0.5e-8
1e-8
1.5e-8
2e-8
2.5e-8
3e-8
3.5e-8
0.5e8
1e8
1.5e8
2e8
0.1e8
0.2e8
0.3e8
0.4e8
0.6e8

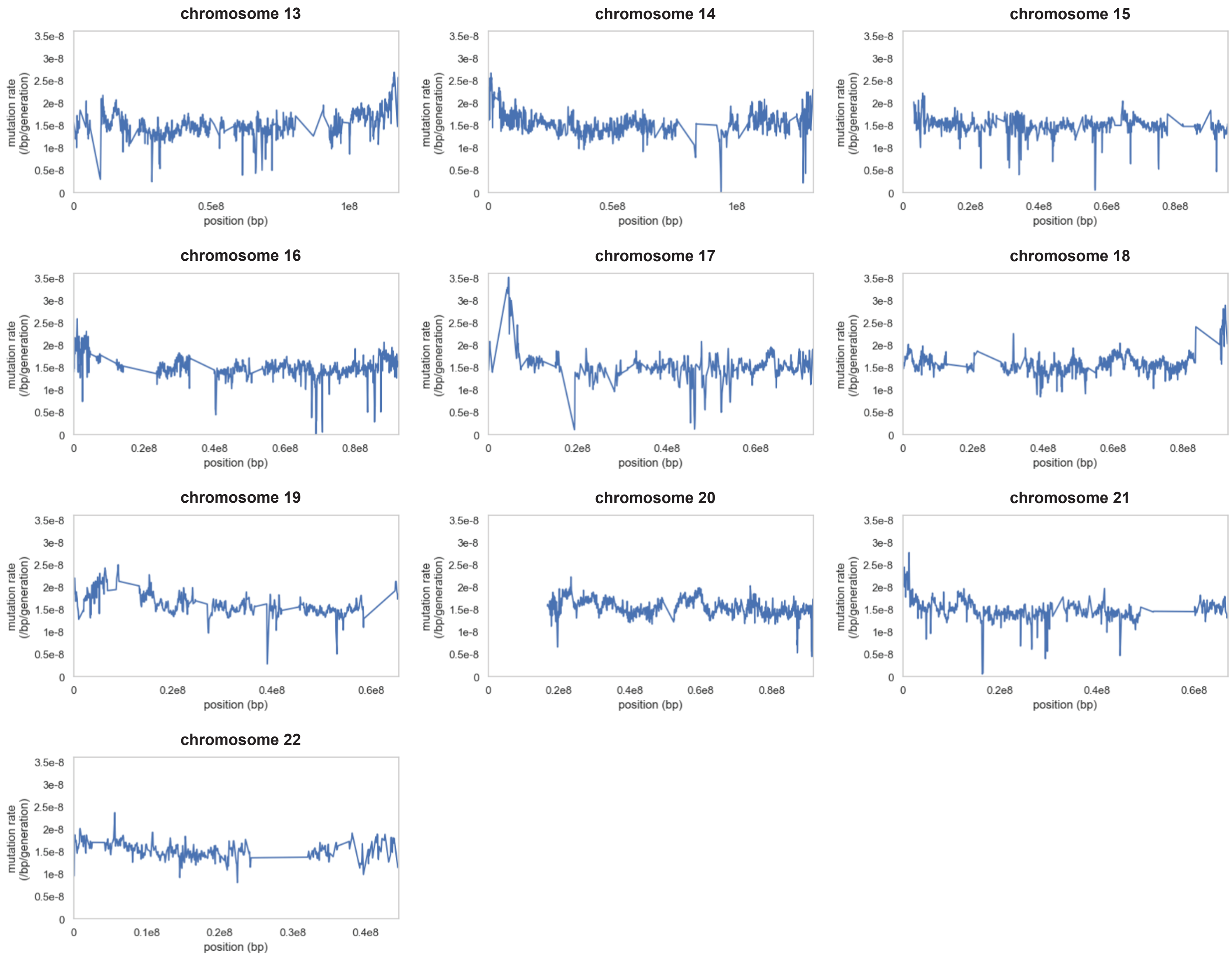


**Supplementary Figure S2.** Per-site per-generation neutral mutation rates across the autosomes for genomic windows of size 100 kb, with a 50 kb step size, assuming a *P. cupreus*–*H. sapiens* divergence time of 33 mya and a generation time of 9 years.

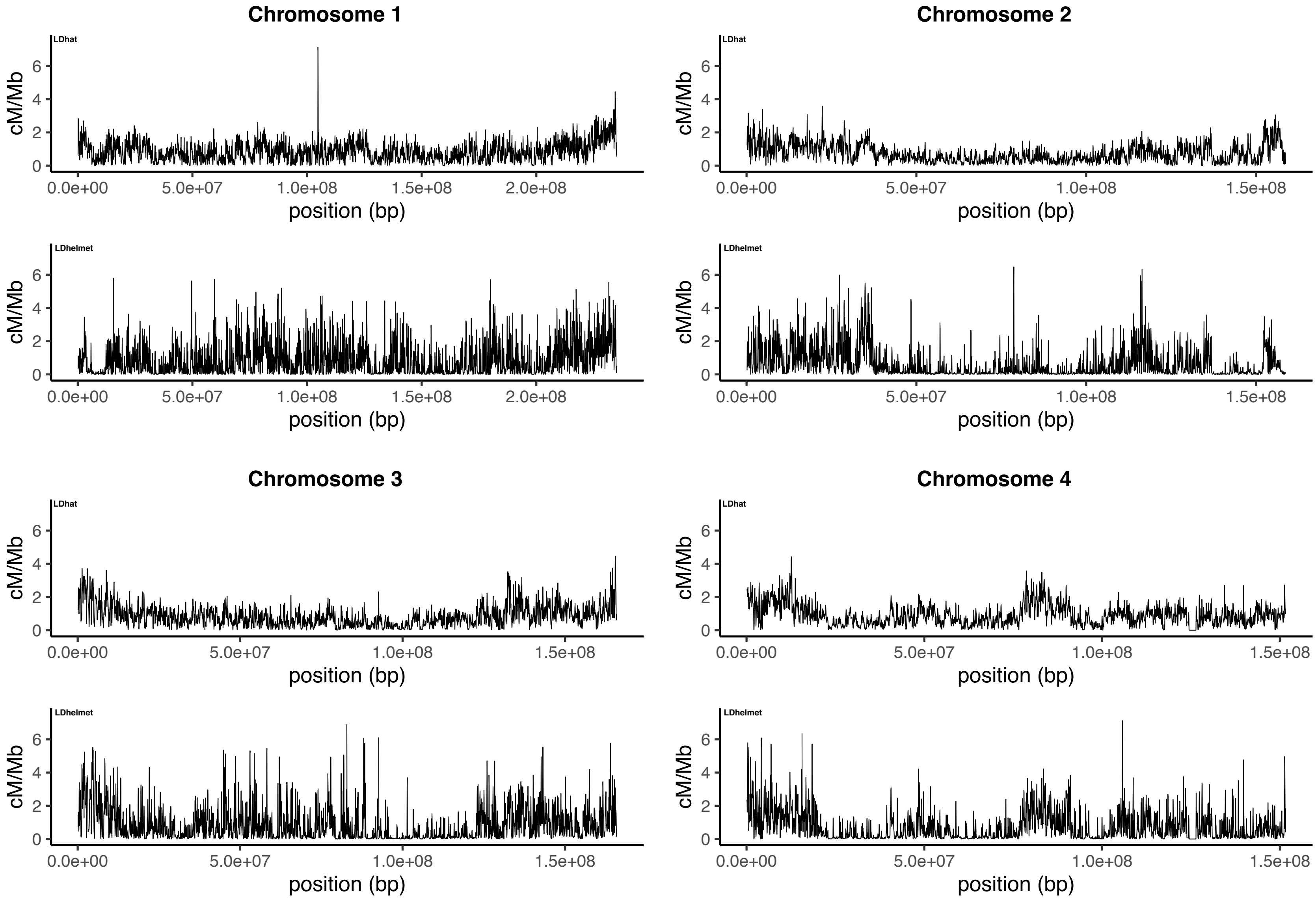

Chromosome 1
Chromosome 2
Chromosome 3
Chromosome 4
LDhat
LDhelmet
cM/Mb
position (bp)
0
2
4
6
0.0e+00
5.0e+07
1.0e+08
1.5e+08
2.0e+08

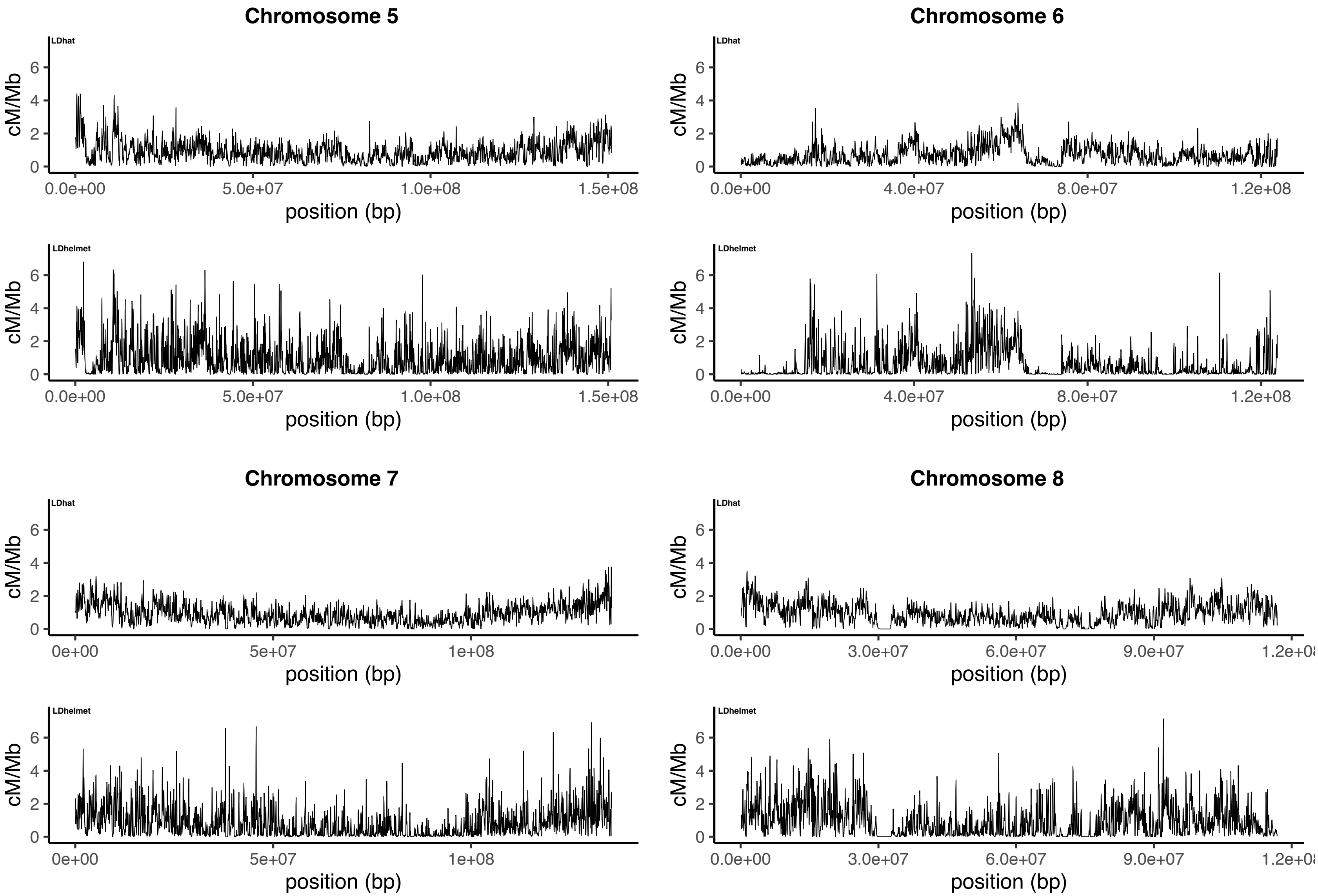

Chromosome 5
LDhat
cM/Mb
0
2
4
6
0.0e+00
5.0e+07
1.0e+08
1.5e+08
position (bp)
LDhelmet
Chromosome 6
LDhat
0.0e+00
4.0e+07
8.0e+07
1.2e+08
LDhelmet
Chromosome 7
LDhat
0e+00
5e+07
1e+08
LDhelmet
Chromosome 8
LDhat
0.0e+00
3.0e+07
6.0e+07
9.0e+07
LDhelmet

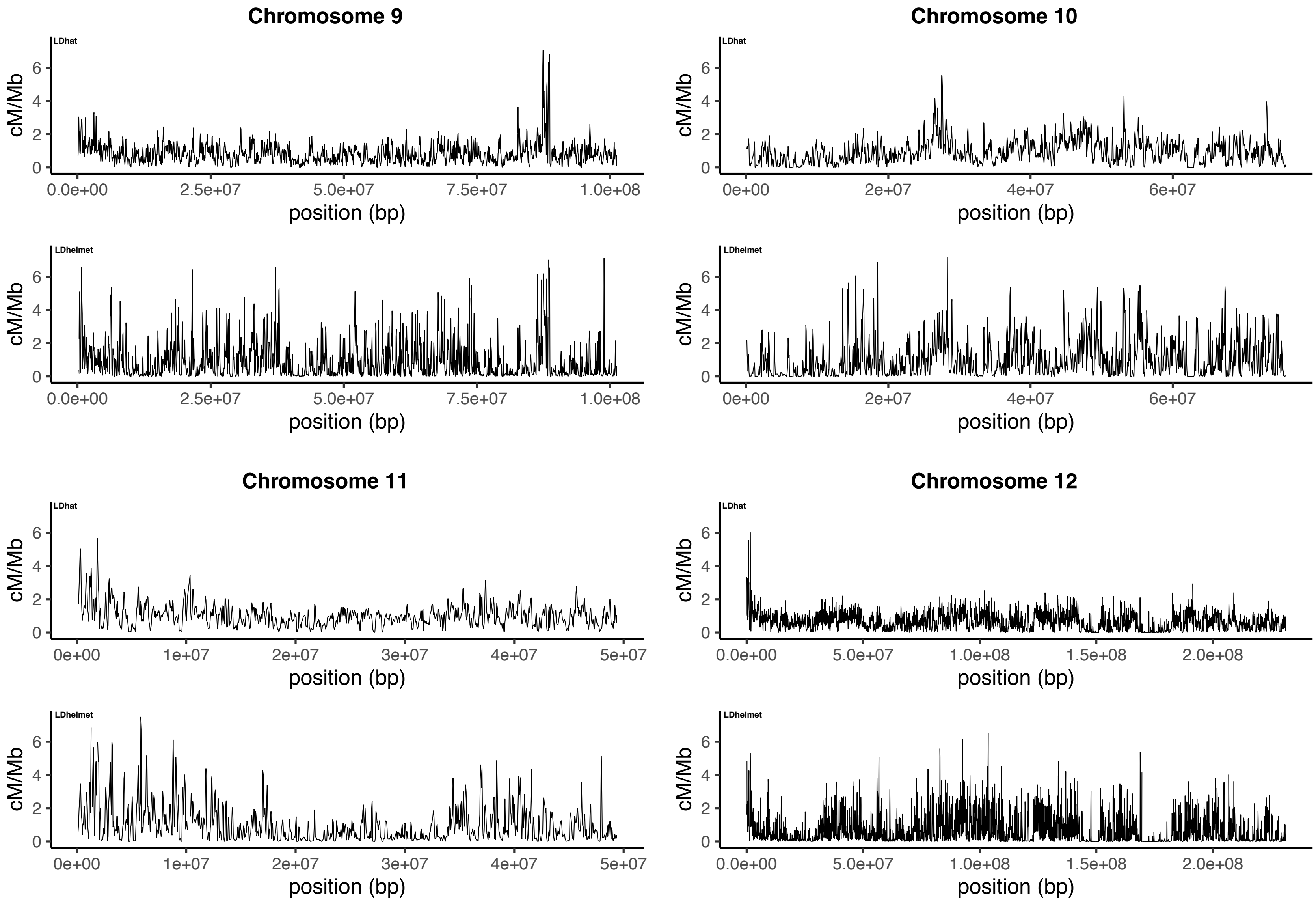

Chromosome 9
LDhat
cM/Mb
0
2
4
6
0.0e+00
2.5e+07
5.0e+07
7.5e+07
1.0e+08
position (bp)
LDhelmet
cM/Mb
position (bp)
Chromosome 10
LDhat
cM/Mb
0e+00
2e+07
4e+07
6e+07
position (bp)
LDhelmet
cM/Mb
position (bp)
Chromosome 11
LDhat
cM/Mb
0e+00
1e+07
2e+07
3e+07
4e+07
5e+07
position (bp)
LDhelmet
cM/Mb
position (bp)
Chromosome 12
LDhat
cM/Mb
0.0e+00
5.0e+07
1.0e+08
1.5e+08
2.0e+08
position (bp)
LDhelmet
cM/Mb
position (bp)

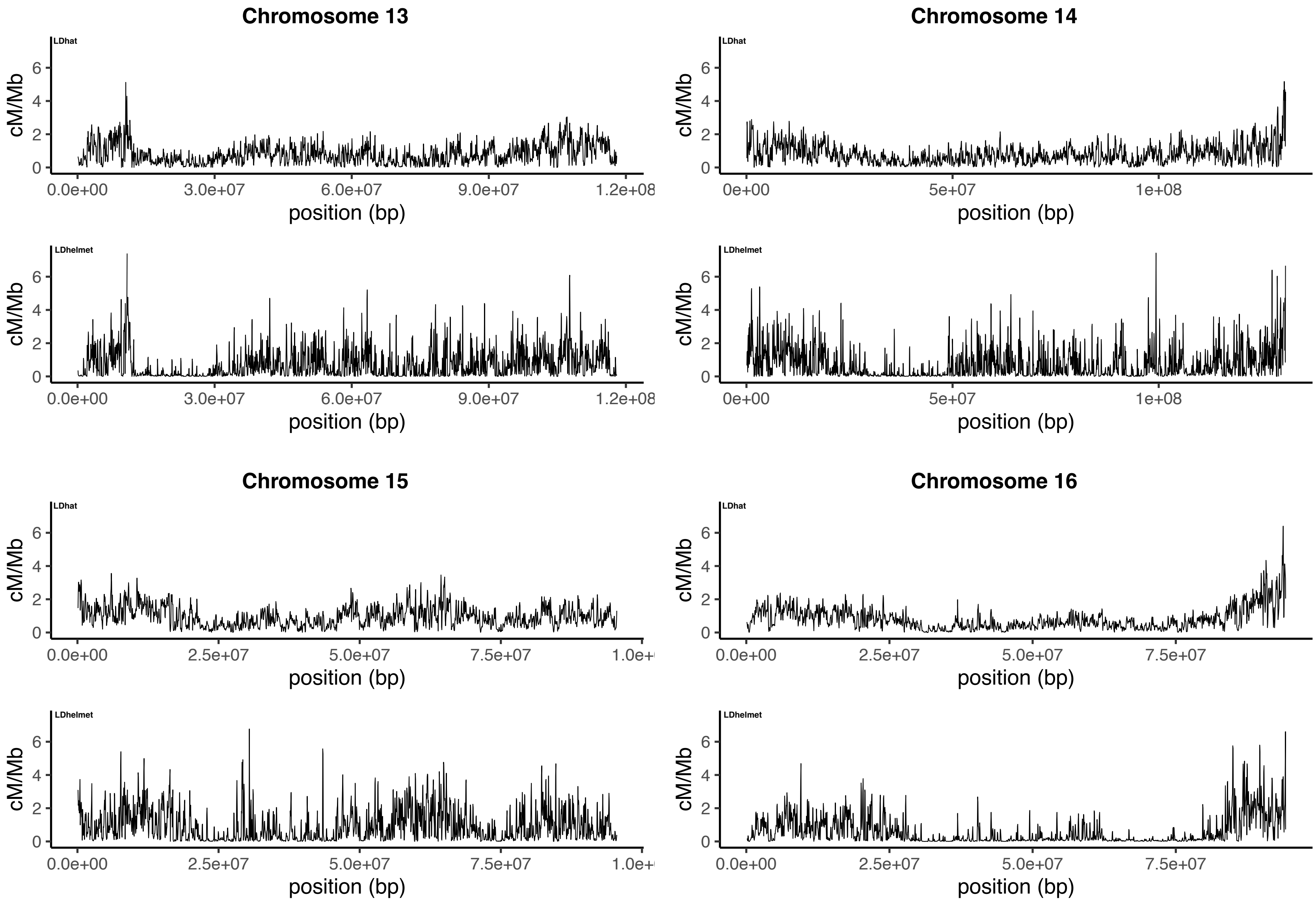
Chromosome 13
LDhat
cM/Mb
0
2
4
6
0.0e+00
3.0e+07
6.0e+07
9.0e+07
1.2e+08
position (bp)
LDhelmet
Chromosome 14
0e+00
5e+07
1e+08
Chromosome 15
2.5e+07
5.0e+07
7.5e+07
1.0e+
Chromosome 16

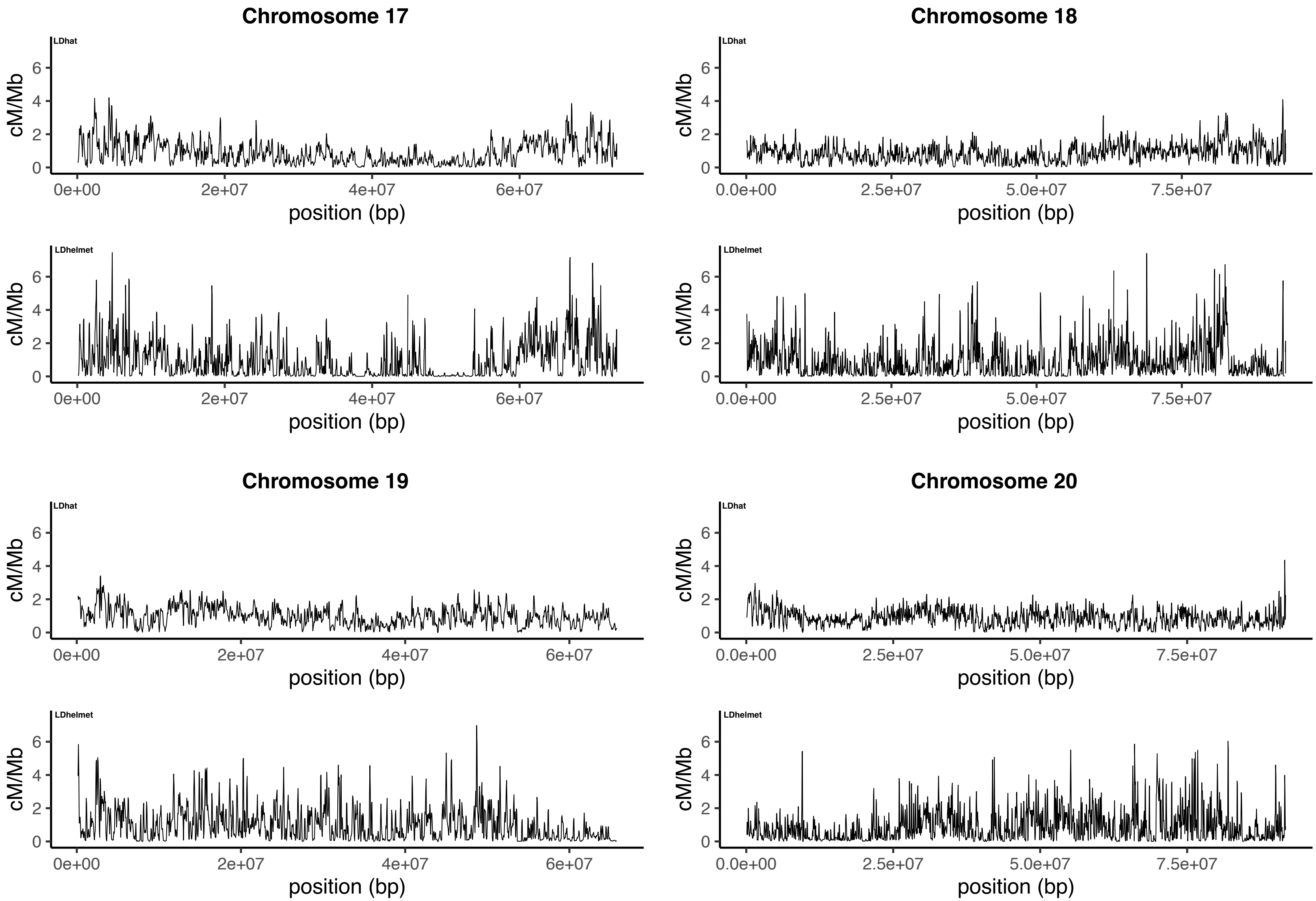
Chromosome 17
LDhat
cM/Mb
0 2 4 6
0e+00 2e+07 4e+07 6e+07
position (bp)
LDhelmet
cM/Mb
0 2 4 6
0e+00 2e+07 4e+07 6e+07
position (bp)
Chromosome 18
LDhat
cM/Mb
0 2 4 6
0.0e+00 2.5e+07 5.0e+07 7.5e+07
position (bp)
LDhelmet
cM/Mb
0 2 4 6
0.0e+00 2.5e+07 5.0e+07 7.5e+07
position (bp)
Chromosome 19
LDhat
cM/Mb
0 2 4 6
0e+00 2e+07 4e+07 6e+07
position (bp)
LDhelmet
cM/Mb
0 2 4 6
0e+00 2e+07 4e+07 6e+07
position (bp)
Chromosome 20
LDhat
cM/Mb
0 2 4 6
0.0e+00 2.5e+07 5.0e+07 7.5e+07
position (bp)
LDhelmet
cM/Mb
0 2 4 6
0.0e+00 2.5e+07 5.0e+07 7.5e+07
position (bp)

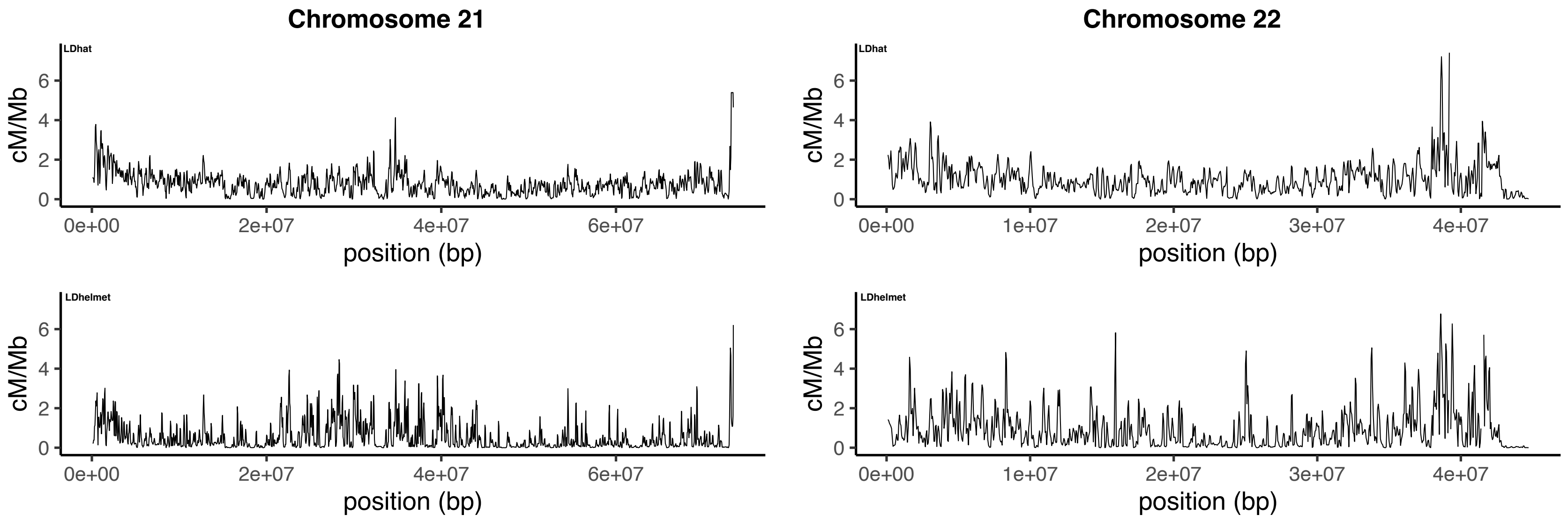


**Supplementary Figure S3.** Recombination rates inferred using LDhat (top) and LDhelmet (bottom) across the autosomes for genomic windows of size 1 Mb, with a 500 kb step size. Results for LDhat and LDhelmet are shown for a block penalty of 5.

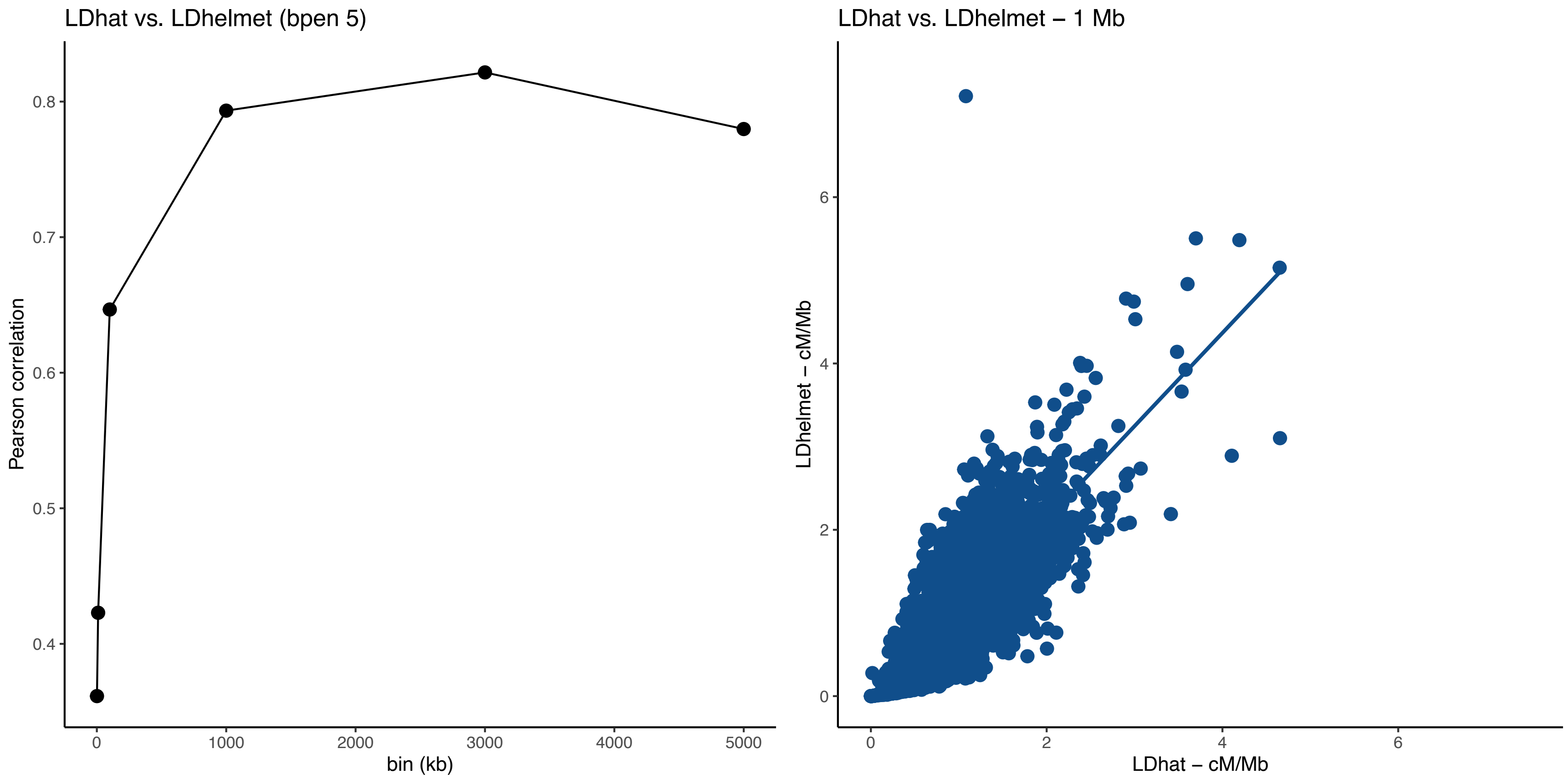


**Supplementary Figure S4. Left panel:** Pearson correlation between LDhat and LDhelmet recombination rate estimates calculated across non-overlapping genomic window of increasing size. **Right panel:** Pearson correlation between LDhat and LDhelmet recombination rate estimates at the 1 Mb scale. Results for LDhat and LDhelmet are shown for a block penalty of 5.
